# Pervasive conditional selection of driver mutations and modular epistasis networks in cancer

**DOI:** 10.1101/2022.01.10.475617

**Authors:** Jaime Iranzo, George Gruenhagen, Jorge Calle-Espinosa, Eugene V. Koonin

## Abstract

Cancer driver mutations often display mutual exclusion or co-occurrence, underscoring the key role of epistasis in carcinogenesis. However, estimating the magnitude of epistasis and quantifying its effect on tumor evolution remains a challenge. We developed a method to quantify *COnditional SELection on the Excess of Nonsynonymous Substitutions* (*Coselens*) in cancer genes. *Coselens* infers the number of drivers per gene in different partitions of a cancer genomics dataset using covariance-based mutation models and determines whether coding mutations in a gene affect selection for drivers in any other gene. Using *Coselens*, we identified 296 conditionally selected gene pairs across 16 cancer types in the TCGA dataset. Conditional selection affects 25-50% of driver substitutions in tumors with >2 drivers. Conditionally co-selected genes form modular networks, whose structures challenge the traditional interpretation of within-pathway mutual exclusivity and across-pathway synergy, suggesting a more complex scenario, where gene-specific across-pathway epistasis shapes differentiated cancer subtypes.

## Introduction

Cancer is a result of somatic evolution, during which cells accumulate driver mutations (drivers for short) that enable them to proliferate, bypassing cell cycle checkpoints and avoiding senescence, invade the surrounding tissues, escape the immune system, and ultimately disseminate to distant body locations(Hanahan and Weinberg, 2000; Stratton et al., 2009). By virtue of increasing the growth and survival rates of (pre)cancer cells, drivers are positively selected and can reach high prevalence within the tumor although not necessarily fixed (Garraway and Lander, 2013; Greaves and Maley, 2012; Nik-Zainal et al., 2012; Vogelstein et al., 2013; Williams et al., 2018). Concomitantly, (pre)cancer cells acquire passenger mutations which, despite not contributing to cancer initiation or progression, can spread by hitchhiking on drivers and typically are the dominant component of cancer mutational landscapes(Martincorena et al., 2017; Weghorn and Sunyaev, 2017).

The precise set of drivers acquired by a tumor determines some of its fundamental properties, such as metastatic potential and sensitivity to targeted therapies(Senft et al., 2017). Some of these properties can be related to the presence or absence of a single driver, whereas others result from the combination of multiple drivers that interact synergistically or interfere with each other, critically affecting tumor initiation, progression, and clinical outcome(Ashworth et al., 2011; Mina et al., 2017; Zhang et al., 2017). Two well-studied examples involve the synergy between activating mutations in oncogene *RAS* and inactivation of the tumor suppressor *ATM*, and the synthetic lethality of *EGFR* and *KRAS* double mutants in lung and colon cancers. In the first case, inactivation of ATM allows tumor cells to overcome oncogene-induced senescence associated with RAS activation, and therefore exacerbates cell proliferation(Di Micco et al., 2006). In the second case, oncogenic mutations in *KRAS* make EGFR-mutant tumors resistant to therapies that target EGFR, but the synergistic deleterious effect of *KRAS* and *EGFR* mutations makes such escape clones inviable in the absence of the drug(Misale et al., 2014).

Epistatic interactions, such as those mentioned above, have the potential to substantially affect cancer somatic evolution by making selection on a driver conditional on the presence or absence of another driver (Wilkins et al., 2018). Epistasis results in patterns of co-occurrence when synergy between drivers enhances positive selection or mutual exclusivity when redundancy or antagonism diminish selection on the double mutant. Epistasis-driven conditional selection implies that past driver mutations affect the fitness of subsequent driver mutations and, with that, the evolutionary trajectory of the tumor.

Searching for mutually exclusive mutations is a main focus of research in cancer genomics because such findings can lead to identification of synthetic lethal pairs and cancer-associated pathways, which are potential therapeutic targets(Babur et al., 2015; Ciriello et al., 2012; Jerby-Arnon et al., 2014; Kim et al., 2017; Leiserson et al., 2015; Matlak and Szczurek, 2017; Mina et al., 2017; Srihari et al., 2015; Wappett et al., 2016). However, because co-occurrence and mutual exclusivity can also result from systematic biases in tumor mutational load, their presence cannot be taken as unequivocal evidence of epistasis(van de Haar et al., 2019). Furthermore, the statistical frameworks commonly used to test for mutation co-occurrence and mutual exclusivity fail to provide quantitative estimates of biologically meaningful effect sizes, such as the extent to which epistasis modifies the fitness of a mutant clone and the subsequent probability that a mutation becomes a driver(Cannataro et al., 2018).

Obtaining quantitative estimates of epistatic effects in a systematic and direct manner is extremely challenging, although a recent study has approached this goal by conducting combinatorial RNAi screening in cell culture followed by image-based phenotyping of growth-related traits(Wang et al., 2014). An alternative approach is to quantify the contribution of epistasis to cancer development indirectly, through its effect on the conditional selection of driver mutations. This approach, which takes advantage of the recent surge in cancer genomic data, has the fundamental advantage of not relying on ad-hoc (*in vitro*) proxies for *in vivo* tumor fitness. A major difficulty, however, is that quantification of conditional selection requires disentangling the contributions of mutation and selection to cancer genomic landscapes, a challenging task that has only recently begun to be addressed by adapting tools from evolutionary genetics for cancer genomics(Cannataro et al., 2018; Martincorena et al., 2017; Persi et al., 2018; Weghorn and Sunyaev, 2017).

Along these lines, we developed a method, Coselens (COnditional SELection on the Excess of Nonsynonymous Substitutions), that compares the mean number of drivers per tumor in a gene depending on whether a second gene harbors non-synonymous mutations, returning an estimate for the magnitude of conditional selection, while controlling for differences in mutational load and spectrum. The principal advantages of Coselens over previously developed methods for analysis of epistasis in cancer is that this method rigorously controls for the biases introduced by cancer subtypes and explicitly estimates the size of epistatic effects. Application of Coselens to somatic mutation data from more than 10,000 cancer exomes led to the identification of a complex network of cancer type-specific epistasis, shedding light on the underlying genetic constraints that shape cancer development.

## Results and Discussion

### Detection of pairwise conditional selection among cancer genes

Driver mutations have been the subject of intense research in recent years, leading to multiple definitions and strategies for driver detection. In this work, we adopted an evolutionary genetics approach, defining drivers as mutations that provide (pre)cancer cells with a selective advantage, and therefore are subject to positive selection. Because purifying selection is typically negligible in non-hypermutated tumors (Martincorena et al., 2017; Persi et al., 2018), the number of positively selected mutations (that is, drivers) in a locus can be obtained as the difference between the number of mutations observed in that locus and the number of mutations that would be expected in the absence of selection. The main challenge of this approach is to infer the null mutation model under the assumption of neutrality, which we did by combining information from synonymous mutations and genomic covariates. This method accounts for the possibility that the same mutation does not necessarily act as driver in different types of tumors and provides a quantitative estimate of the probability that a particular mutation, if observed, is indeed a driver. Although this definition applies, in principle, to any kind of mutation, we focused on nonsynonymous and truncating substitutions as driver candidates because those are the mutations for which the most reliable null models can be built.

To investigate the role of conditional selection in cancer progression, we quantified the dependency of the number of driver substitutions in a cancer gene (the “query” gene) on the presence or absence of coding mutations in a second gene (the “split” gene). For each cancer type in The Cancer Genome Atlas (TCGA), we first identified the set of query genes by pooling all somatic mutations from the same cancer type and searching for genes for which the dN/dS indicated significant levels of positive selection (Supplementary Table 1). Query genes with coding mutations (nonsynonymous substitutions, truncations, and small indels) in >25 samples were additionally considered as split genes. Then, for all split-query gene pairs, we classified tumor samples into those that did and did not harbor coding mutations in the split gene and applied our evolutionary genetics-inspired method *Coselens* to estimate and compare the number of driver substitutions in the query gene in each of the groups (Figure 1a). In the first step, *Coselens* fits a mutational model to each group of samples and calculates the mean number of nonsynonymous and truncating substitutions that would be expected in the query gene in the absence of selection. Then, it calculates the number of drivers as the difference between the number of observed and expected mutations (see Methods for details). Finally, *Coselens* implements a likelihood ratio test to assess whether the number of driver events significantly differs between samples that do and do not harbor coding mutations in the split gene.

**Figure 1:**
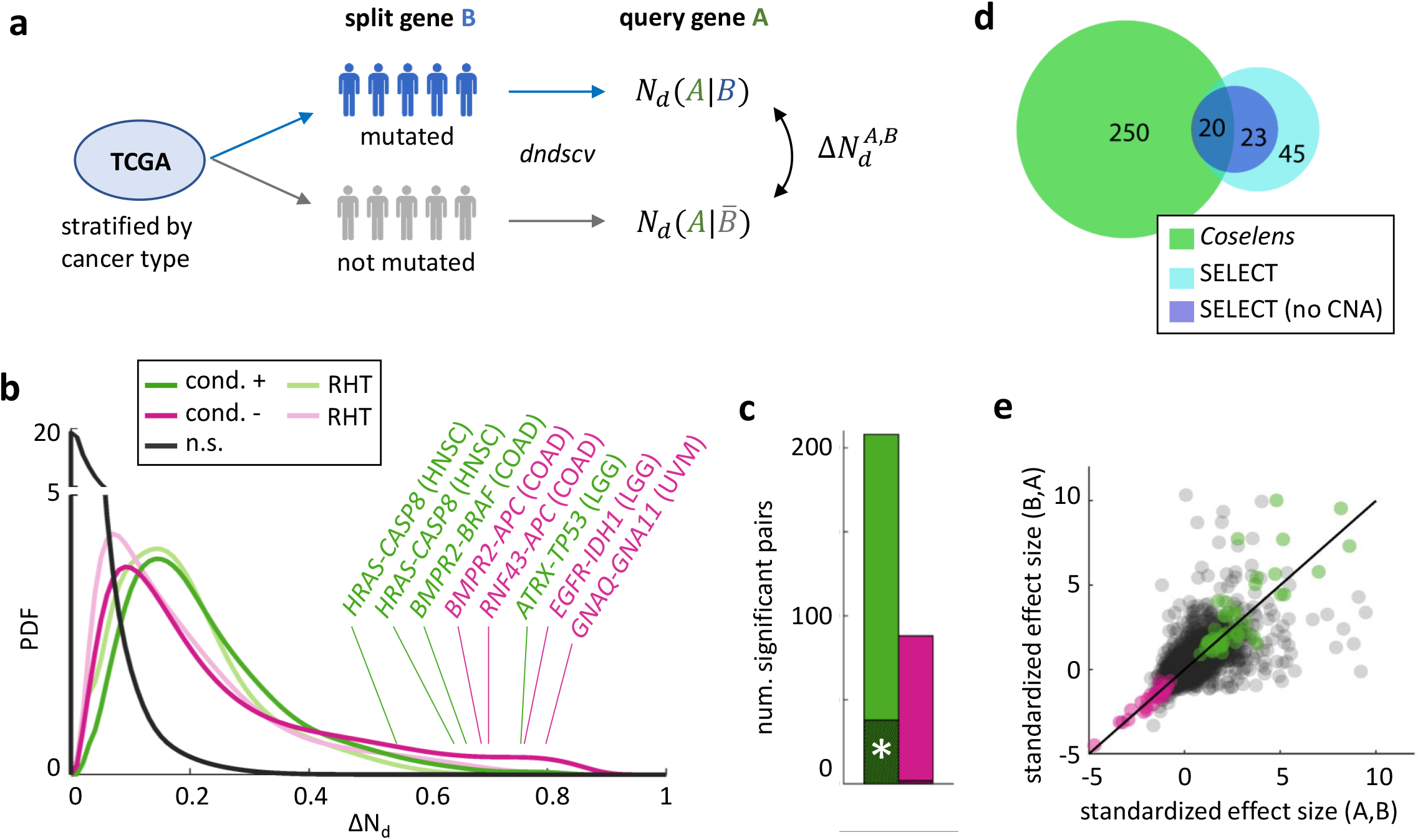
Inference of pairwise conditional selection in cancer genes. a) Overview of the *Coselens* method. The effect size Δ*N*_*d*_ is the difference in the frequency of driver substitutions in a given gene (the query gene) between tumors that do and do not harbor coding mutations in a second gene (the split gene). b) Effect size distributions for statistically significant and not-significant conditional gene pairs. Negative effect sizes are represented by their absolute values. RHT, statistically significant pairs under Restricted Hypothesis Testing of reciprocal interactions. c) Number of gene pairs subject to significant conditional selection (FDR ≤ 0.05), classified by the sign of their interaction. The shaded green bar (*) indicates a small subset of significant positive pairs whose conditional mutation patterns can be explained by differences in mutation availability. d) Comparison of *Coselens* and SELECT(Mina et al., 2017) methods for the detection of conditionally selected gene pairs. CNA: copy number alterations. e) Comparison of the standardized effect sizes that result from exchanging the query gene by the split gene (an outlier at [18.4, 8.7] is not shown). The symmetry of standardized effect sizes implies that passenger mutations in the split gene do not substantially affect the estimates of conditional selection acting on the query gene.

Our analysis identified 296 gene pairs subject to significant conditional selection (FDR ≤ 5%) in at least one cancer type (Supplementary Table 2). Of these, 262 were deemed significant by the standard workflow and the rest were identified through restricted hypothesis testing of reciprocal conditional selection (that is, by testing the reverse pairs, in which the split and query genes were swapped). The effect size distribution of significant pairs identified through restricted hypothesis testing closely resembles that of significant pairs detected through the more stringent, standard workflow (Figure 1b), suggesting that the former represent genuine cases of conditional selection. Overall, conditional selection affects 112 genes across 16 cancer types. In 70% of the conditionally selected gene pairs, coding mutations in the split gene increase the probability of finding driver mutations in the query gene; henceforth, we refer to these conditional pairs as positive. In the remaining 30%, coding mutations in the split gene decrease the probability to find driver mutations in the query gene, and henceforth, we refer to these conditional pairs as negative (Figure 1c).

The magnitude of conditional selection varies among gene pairs, with coding mutations in the split gene typically associated with an increase or decrease of at least 0.15 driver substitutions per query gene per tumor (Figure 1b). In some extreme cases, conditional selection results in almost complete presence or absence of a driver mutation in the query gene. The pair of closely related oncogenes *GNAQ* and *GNA11* is a good example of such extreme conditionality: driver substitutions in either GNAQ or GNA11 are found in 80-90% of uveal melanomas. However, mutations in these genes are mutually exclusive(Decatur et al., 2016; Van Raamsdonk et al., 2010), so that the absence of mutations in one implies, with high probability, the presence of driver mutations in the other. On the other extreme, the pair formed by *TP53* and *ATRX* shows strong, mutual positive conditionality in low-grade gliomas, such that driver substitutions in *ATRX* appear almost exclusively in mutated *TP53* backgrounds(Liu et al., 2012).

We compared the results of *Coselens* with those obtained using a mutual information approach (*SELECT*), which was also developed to identify conditional selection in cancer(Mina et al., 2017). The comparison indicates that *Coselens* detects many more conditionally selected gene pairs than *SELECT*, but the two approaches seem to capture complementary aspects of conditional selection in cancer genomes (Figure 1d). By leveraging information from copy number alterations (CNA), *SELECT* can detect conditional selection in genes whose role in cancer depends on amplification and/or deletion, albeit at the cost of not providing easily interpretable effect sizes. In contrast, *Coselens* proves substantially more powerful at detecting and quantifying conditional selection among cancer genes that harbor driver substitutions, with 7-and 4-fold increases in the number of positive and negative conditional pairs compared to *SELECT* (Supplementary Figure 1). Such higher detection power can be partly explained by *Coselens*’ ability to analyze genes that fail some of the pre-filtering criteria applied by *SELECT* (this is the case for 79 of the 250 *Coselens*-exclusive pairs). In turn, 17 of the 23 non-CNA, *SELECT*-exclusive pairs involved genes that did not show evidence of positive selection as per their *dN/dS* values.

### Mutational load does not substantially confound *Coselens*’ estimates of conditional selection

Previous studies have noted that detection of mutual exclusivity can be confounded by intrinsic differences in the frequency and spectrum of mutations across cancer subtypes(Kim et al., 2017; van de Haar et al., 2019). To minimize this risk, *Coselens* independently fits two baseline mutation models, one for samples with coding mutations in the split gene and another for samples without such mutations. The use of subset-specific models for the expected number of substitutions helps prevent spurious correlations resulting from across-subset variations in mutational signatures. However, differences in mutational load can also affect the detection of conditional selection indirectly, in cases when mutation availability is a limiting factor in tumor evolution. If differences in mutational load significantly affect the availability of mutations on which selection can act, then, associations between rare cancer genes could reflect patterns of differential mutation availability rather than conditional selection. To account for that possibility, we adopted two alternative correction methods: one applies sampling theory to assess the likelihood that each significant gene pair can be explained by differences in mutation availability, and the other uses regression to identify biases in the number of significant pairs that correlate with mutation load (see Methods). The results obtained with both methods were in close agreement, leading to the conclusion that >80% of conditional gene pairs could not be explained by differential mutation availability (Figure 1c).

We also investigated whether nonsynonymous passenger mutations in split genes quantitatively affected the inference of conditional selection. Because conditional selection is inferred by partitioning tumors into those with and without coding mutations in one of the genes, the inclusion of non-driver mutants in the former set could lead to underestimating the real magnitude of the association between driver mutations. To test whether estimates of conditional selection are indeed affected by passenger coding mutations in the split gene, we used Bayes theorem to derive an equivalence expression that relates the standardized effect size of conditional selection upon swapping the split and query genes (see Methods). This equivalence should hold exactly in the case when conditional selection results from epistasis between driver mutations, but deviations are expected to appear if the inference of conditional selection is biased by passenger coding mutations in the split gene. Following this rationale, we compared the standardized effect sizes of conditional selection in swapped gene pairs (Figure 1e). The close agreement between empirical estimates and theoretical predictions (intraclass correlation = 0.88, N = 78, p<1e-4) confirms that estimates of coding-to-driver conditional selection are good proxies for the magnitude of driver-to-driver epistasis.

### Functional association and classification of epistasis in conditionally selected gene pairs

To better characterize the mechanisms underlying conditional selection, we searched the STRING database for functional and physical interactions between the split- and query-gene products. Based on that search, we calculated an interaction enrichment ratio that quantifies, for any subset of gene pairs, the enrichment in predicted interactions of different types with respect to the set of all tested gene pairs. Given that STRING compiles extensive information on various types of association between genes and the encoded proteins, including functional data extracted from the literature, experimental evidence of physical interactions, and coexpression across a broad diversity of organisms (Szklarczyk et al., 2021), we considered it the best available source of background knowledge of gene-gene interactions. We found that enrichment in interactions increases with the increase of the effect size of conditional selection (Figure 2a). Pairs subject to significant conditional selection encompass more interactions than expected by chance, with a 1.5-fold enrichment (95% CI 1.2-2.0) in the case of positive conditional pairs and a 4.2-fold enrichment (95% CI 2.7-6.6) in the case of negative conditional pairs (Figure 2b). Regarding the specific nature of interactions, no significant differences were found between functional association and physical binding, although the latter seem to be slightly more prevalent among the negative conditional pairs.

**Figure 2:**
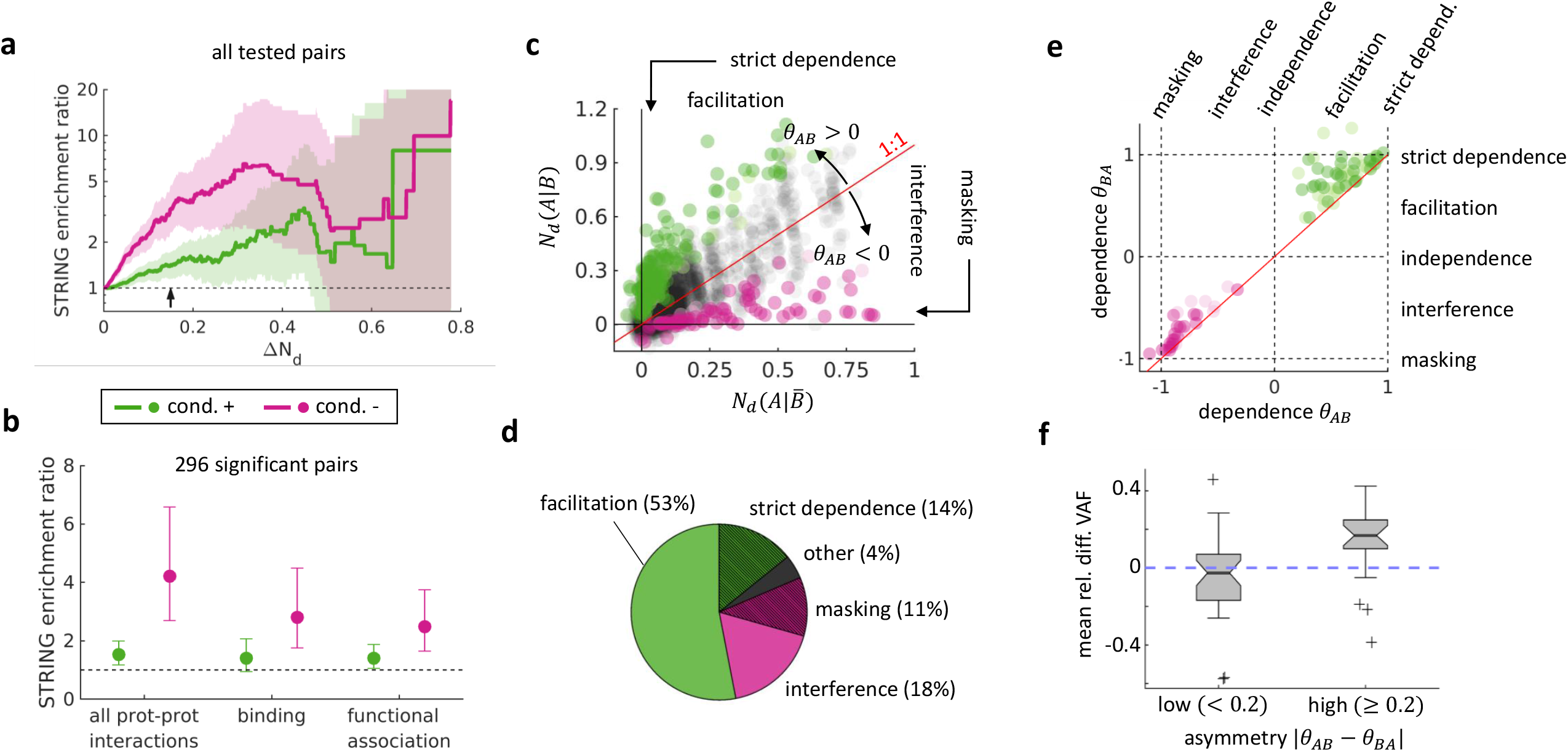
Functional association and epistasis classification of conditionally selected gene pairs. a) Protein-protein interactions between gene products of the same conditional pair. The STRING enrichment ratio (y axis) was calculated using the formula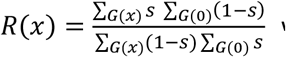, where *s* is the STRING score (rescaled in the range 0-1) and *G*(*x*) is the set of conditional gene pairs such that Δ*N*_*d*_ ≥ *x* (*x* axis). *R*(*x*), which is formally analogous to a risk ratio for continuous response variables, compares gene pairs above a given intensity of conditional selection with a reference base value calculated using all possible gene pairs. The small arrow at Δ*N*_*d*_ = 0.15 indicates the effect size of conditional selection at which most interactions become statistically significant. Shaded regions correspond to 95% confidence intervals. The noisy values at Δ*N*_*d*_ > 0.4 are due to the small number of gene pairs that reach such high levels of conditional selection. b) Enrichment in different types of protein-protein interactions, calculated for the 296 gene pairs subject to significant conditional selection (whiskers indicate 95% confidence intervals). c) Epistasis is quantified attending to the frequency of driver substitutions in the query gene when the split gene does not harbor coding mutations (x axis) and when it does (y axis). In the absence of epistasis, the frequency of drivers in the query gene is not affected by the mutation status of the split gene, resulting in a 1:1 trend. Facilitation and interference manifest as positive and negative deviations from that trend and their magnitude is quantified by the angle *θ* (normalized by a factor *π*/4). Strict dependence and masking correspond to the extreme cases *x* = 0 (*θ* = 1) and *y* = 0 (*θ* = −1), respectively. None of the negative data points in the figure significatively deviates from 0. d) Relative abundances of different classes of epistasis among 296 significantly conditional gene pairs. e) Asymmetry of epistasis. Each data point corresponds to a statistically significant conditional gene pair whose reversed pair (with split and query gene roles exchanged) is also significant. The diagonal represents purely symmetric epistasis, whereby both genes equally affect selection on each other. f) Mean relative difference in the variant allele frequencies for coding mutations in genes subject to reciprocal positive epistasis. Gene pairs were grouped based on the asymmetry of epistasis (low asymmetry if |*θ*_*AB*_ − *θ*_*BA*_| < 0.2; high asymmetry if |*θ*_*AB*_ − *θ*_*BA*_| ≥ 0.2). Boxes comprise the 25th-75th percentiles; thick lines and notches represent the median and its 95% confidence interval, respectively; whiskers extend to the last data point within 1.5 times the interquartile range; outliers beyond that range are represented as with the symbol +.

Epistasis between cancer genes can be classified attending to the sign and magnitude of the conditional selection on driver substitutions. We identified 4 major classes of epistasis (Figure 2c): *strict dependence*, when driver (that is, positively selected) substitutions in the query gene are only observed in the presence of coding mutations in the split gene; *facilitation*, when driver substitutions are more likely to occur when the split gene is mutated; *interference*, when driver substitutions are less likely to occur when the split gene is mutated; and *masking*, when mutations in the split gene completely abolish the effect of driver substitutions in the query gene. Although strict dependence and masking appear as the extremes in the continuum that goes from facilitation to independence to interference, they can be distinguished by testing the null hypothesis that there is no significant selection on nonsynonymous mutations. By far, the most prevalent form of conditional selection is facilitation, which was identified in more than half of the significant gene pairs (Figure 2d). Focusing on the extreme classes, masking represents almost 40% of negative epistatic interactions (11% of the total), whereas strict dependence only represents 20% of positive interactions (14% of the total).

Positive and negative epistasis differ in the degree of mutual interdependency between the two genes of the same pair (Figure 2e). Negative epistasis tends to be symmetrical and often involves mutual masking. This is consistent with negative epistasis arising from complete functional redundancy although it could also reflect the existence of multiple, incompatible mutational paths to cancer (see below). On the other side, the spectrum of positive epistasis is broader and combines cases of strict interdependence and asymmetric facilitation. A biologically plausible interpretation of asymmetric facilitation is that driver mutations in a “master” cancer gene open new pathways of tumor development that involve the dependent gene. As a result, mutations in the dependent gene only act as drivers if the master gene carries drivers as well, but not vice versa. Supporting this interpretation, we found that in pairs with moderate and high asymmetry, mutations in the master gene tend to show a higher degree of clonality (mean relative difference in variant allele frequency = 0.13, p = 0.009, 1-sample Student’s t-test, n = 20) and therefore are likely to precede mutations in the dependent gene (Figure 2f). In contrast, no systematic differences in clonality were found between genes that participate in symmetric epistasis. We observed that 6 of the 15 most asymmetric pairs identified in this study involved *PTEN* mutations in endometrial cancers (Supplementary Table 3), which underscores the fundamental role of *PTEN* inactivation as an early driver event in this cancer type(Levine et al., 1998; Zhang and Yu, 2010). Another highly asymmetric pair, observed in bladder and colorectal cancer, consists of *TP53* as the master and *RB1* as the dependent gene. Considering the tumor suppressor role of these genes, the asymmetry of conditional selection implies that, for *RB1* loss to act as a driver, *TP53* must also harbor inactivating driver mutations, whereas *TP53* inactivation can act as a driver on its own. Thus, an intact *TP53* could partially compensate for the loss of *RB1*, but not vice versa.

### Conditional selection across cancer types

Different cancer types broadly vary in the number of gene pairs subject to conditional selection, with the highest numbers found in endometrial and colorectal cancers, followed by stomach and bladder cancers (Figure 3a). To quantify the contribution of conditional selection to cancer somatic evolution, we compared the numbers of driver substitutions per tumor that were affected by conditional selection across cancer types (Figure 3b). We found that positive conditional selection is responsible for the spread of a sizable fraction (25-50%) of all driver substitutions. The relative contributions of interference and masking are generally smaller although in some tumors, such as uveal melanoma, lower glade gliomas, and endometrial cancers, negative epistasis could lead to lack of selection for (and therefore hinder fixation of) >40% potential driver mutations.

**Figure 3:**
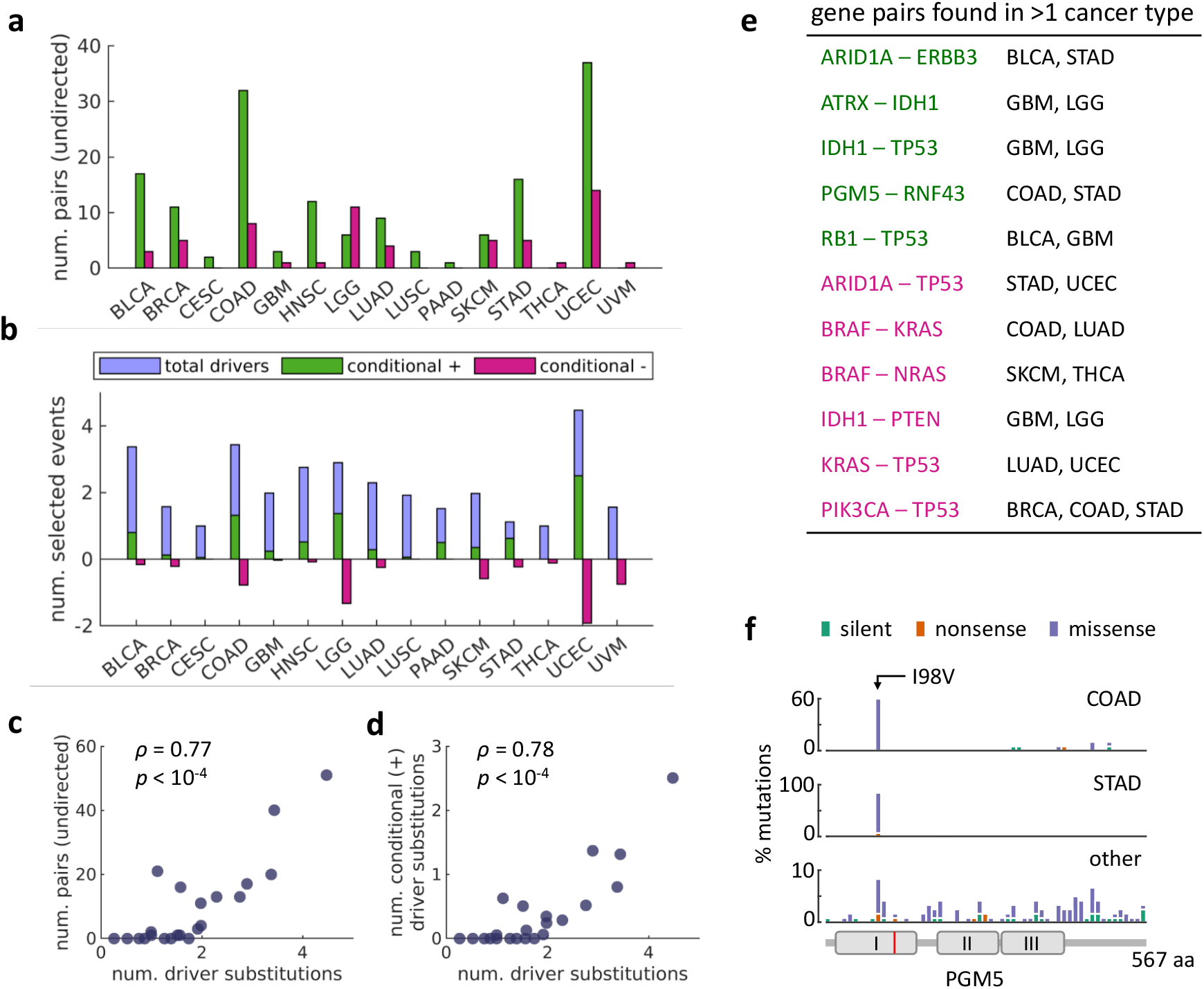
Distribution of conditionally selected gene pairs across cancer types. a) Number of gene pairs affected by significant conditional selection in each cancer type. Gene pairs with reversed split and query genes were counted only once. No significant gene pairs were detected for sarcoma, thymoma, esophageal, renal (clear cell), liver, ovarian, and prostate cancers. b) Number of conditionally selected driver substitutions and total driver substitutions per tumor. Negative values can be interpreted as substitutions that could have been drivers in the absence of epistasis, but did not become established due to lack of selection. c, d) Association between the total number of driver substitutions per tumor (x axis) and (c) the number of significant gene pairs, (d) the number of conditionally selected driver substitutions (y axis). Each data point corresponds to a cancer type (n = 22). These associations remain highly significant after controlling for sample size (number of tumors and total mutations), median mutational load, and number of pairs tested. e) Conditional gene pairs found in multiple cancer types. f) Distribution of mutations along the PGM5 protein in stomach, colorectal, and other cancer types. The cartoon at the bottom indicates the location of alpha-D-phosphohexomutase, alpha/beta/alpha domains I, II, and III, with the predicted active site in red.

Both the number of gene pairs subject to conditional selection and the average number of conditionally-selected driver substitutions strongly correlate with the total number of driver substitutions per tumor (Figure 3c-d). These associations persist when controlling for several possible confounding factors, including the number of samples and mutations per cancer type, median mutational load, and number of gene pairs tested, that might unequally affect our ability to detect conditional selection in different cancer types (Spearman partial correlation coefficients rho = 0.95 (p<10^−8^) for the number of gene pairs and rho = 0.80 (p<10^−4^) for the number of conditional drivers, n = 22). A strong correlation was observed also with the number of non-conditionally-selected driver substitutions (Spearman partial correlation coefficients rho = 0.96 (p<10^−8^) for the number of gene pairs and rho = 0.75 (p<0.001) for the number of conditional drivers, n = 22). Taken together, these observations imply that conditional selection is intrinsic to cancer somatic evolution, its contribution becoming increasingly important as the number of driver mutations increases.

Most gene pairs subject to conditional selection are highly specific to a single cancer type, with only 3% positive and 12% negative conditional pairs found in >1 cancer type (Figure 3d). Such higher promiscuity of negative conditional gene pairs, though small in magnitude, is statistically significant (chi-square = 6.24, p = 0.03, permutation test). Not surprisingly, multi-tissue gene pairs include well-known examples of genes that display mutually exclusive or concurrent mutation patterns in pan-cancer analyses(Kandoth et al., 2013). Among the negative conditional pairs found in multiple cancer types, those involving oncogenes *KRAS*/*NRAS* and *BRAF* are easily ascribed to the functional redundancy in activating the Ras signaling pathway. The mutual exclusivity between *ARID1A* and *TP53* possibly results from functional codependency in suppressing tumor development through transcriptional regulation(Guan et al., 2011; Wu et al., 2014). However, the mechanisms underlying the negative conditionality between mutations in *TP53* and *KRAS* (and also *TP53* and *PIK3CA*) remain unclear and could reflect fundamental, but poorly understood interactions among major pathways involved in cancer development. Positive conditional pairs spanning multiple cancer types include associations among the genes *ATRX, IDH1*, and *TP53*, which are known to harbor concurrent mutations in brain tumors (glioblastoma and lower grade gliomas)(Brennan et al., 2013; Cancer Genome Atlas Research et al., 2015; Liu et al., 2012). Other multi-tissue pairs are *TP53-RB1* (detected in bladder cancer and glioblastoma), *ARID1A-ERBB3* (bladder and stomach cancer), and *PGM5-RNF43* (colorectal and stomach cancer). Interestingly, the gene *PGM5* (encoding phosphoglucomutase-like protein 5) has not been previously reported as a cancer gene, although low *PGM5* expression has been linked to poor survival in liver and colorectal cancers(Jiao et al., 2019; Sun et al., 2019). Almost 75% (35 out of 47) coding mutations in *PGM5* detected in stomach and colorectal cancers correspond to a single nonsynonymous substitution with predicted deleterious effect (SIFT: 0.01 -deleterious; PolyPhen: 0.573 - possibly damaging) affecting position 98 (Ile>Val) of the encoded protein, without loss of the second copy of the gene. In contrast, this substitution only represents 7% (7 out of 96) *PGM5* coding mutations in other cancer types (Figure 3f). Thus, the PGM5 p.I98V mutation is likely to be a specific driver of gastric and colorectal cancers that possibly operates through a gain-of-function mechanism.

Apart from the broadly spread gene pairs described above, the specificity of most conditional pairs for a single cancer type suggests that epistasis is often context-dependent and is conditioned on the physiology of the tumor(Park and Lehner, 2015; Yeang et al., 2008). This context dependency is manifest at two levels: first, cancer genes involved in conditional selection are often specific to one or a small number of cancer (sub)types; second, when the same gene appears in more than one cancer type, it is often involved in different conditional pairs. The first situation is illustrated by *HLA-C*, which we identified as a conditional cancer gene in *BRAF*-mutant colorectal cancer (Figure 4). Conditional selection on *HLA-C* inactivating mutations is probably mediated by the highly immunogenic nature of tumors of the consensus molecular subtype 1 (CMS-1), whereby the loss of MHC-I-mediated antigen presentation confers a selective advantage(Jhunjhunwala et al., 2021). Supporting this interpretation, multiple studies have described driver mutations in *B2M* and *HLA* genes in CMS-1 colorectal cancer and other microsatellite unstable tumors characterized by high levels of CD8+ T cell infiltration(Castro et al., 2019; Kloor et al., 2007; Shukla et al., 2015). The second situation is exemplified by *RNF43* and its different conditionality patterns observed in colorectal, gastric, and endometrial cancers (Figure 4 and Supplementary Figure 2). In agreement with these differences, two distinct mechanisms have been proposed to explain how the loss of *RNF43* contributes to carcinogenesis, with deregulation of Wnt signaling being dominant in the colon and modulation of the DNA damage response playing the principal role in the stomach(Neumeyer et al., 2021; Neumeyer et al., 2019). An even more striking example is presented by conditionally selected mutations in *KRAS* and *TP53*, which are antagonistic in lung and endometrial tumors, but synergistic in pancreatic cancer. For the latter cancer type, recent research has shown that mutant KRAS and p53 interact through the CREB1 protein to promote metastasis(Kim et al., 2021). The pronounced cancer type-specificity of conditional selection among widespread cancer genes highlights the need to integrate functional pleiotropy, cellular environment, and tumor physiology in models of cancer evolution(DeGregori, 2017).

**Figure 4:**
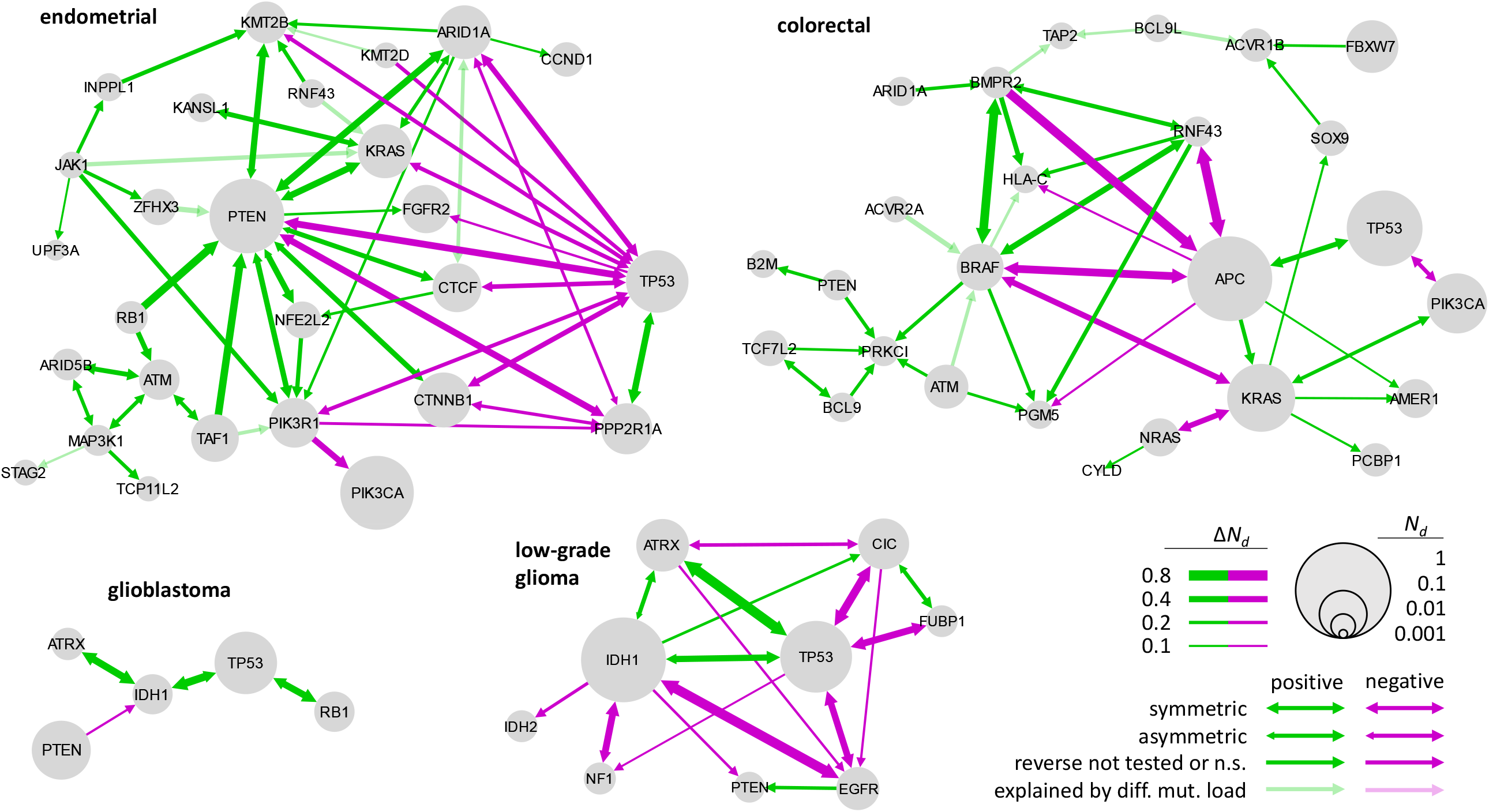
Epistasis networks based on conditional selection for some representative cancer types (see Suppl. Fig. 2 for the remaining cancer types). Nodes represent cancer genes involved in conditional selection; edges denote conditionally selected gene pairs. Cancer genes that are not involved in conditional selection are not represented, although they sometimes harbor a sizeable fraction of driver substitutions. Node size is proportional to the frequency of driver substitutions in a gene; edge width is proportional to the effect size; edge color represents the sign of epistasis; and arrows indicate the degree of symmetry (fully unidirectional arrows indicate lack of statistical significance and/or insufficient sample size for testing the reverse interaction).

### Epistasis networks

To further elucidate how epistasis among somatic mutations affects tumor evolution, we built epistasis networks for each cancer type by connecting the genes that are involved in conditional selection. The most salient feature of such networks is their modular organization (Figure 4). Modularity implies that cancer genes organize into clearly differentiated groups (hereafter modules), such that genes within a module connect to each other through positive epistasis, whereas genes from different modules display negative or no epistasis. Modularity is most evident in cancer types where conditional selection affects a large fraction of drivers, such as low-grade glioma, colorectal, and endometrial cancers.

Inspection of the genes that belong to different modules suggests an association between the modular structure of the epistasis network and the existence of well-defined molecular cancer subtypes. This association is especially clear in low-grade gliomas, where the three main modules encompassing (i) *IDH1, ATRX*, and *TP53*; (ii) *CIC* and *FUBP1*; and (iii) *EGFR* and *PTEN*, correspond, respectively, to three recently proposed molecular subtypes based on the *IDH1* mutation and 1p/19q codeletion status(Cancer Genome Atlas Research et al., 2015; Ceccarelli et al., 2016), whereas negative conditional selection between the genes of the first module and the *NF1* gene is reminiscent of the molecular landscape found in gliomas from patients with neurofibromatosis type 1 and the LGm6 subgroup of sporadic glioma(Ceccarelli et al., 2016; D’Angelo et al., 2019; Verhaak et al., 2010). In glioblastoma, positive conditional selection among *IDH1, ATRX*, and *TP53* defines a module associated with G-CIMP (glioma-CpG island methylator phenotype) tumors from the proneural subtype(Brennan et al., 2013; Noushmehr et al., 2010). To a lesser extent, the high prevalence of mutations in *BRAF* suggests a connection between consensus molecular subtype 1 of colorectal cancer and a module involving *BRAF, BMPR2, RNF43* and *HLA-C* (Guinney et al., 2015). Similarly, in endometrial cancer, the detection of positive conditional selection between *TP53* and *PPP2R1A* and the antagonism between these two genes and *PTEN* are consistent with the mutational landscape of serous endometrial tumors of the “copy number-high” molecular subtype(Cancer Genome Atlas Research et al., 2013; McConechy et al., 2011).

A possible explanation for the association between conditionally selected gene sets and cancer subtypes could be that the latter are a confounding factor for the inference of conditional selection. More precisely, because all samples from the same cancer type are pooled for the analysis, groups of genes that are preferentially mutated in a single subtype but not in the rest could appear as being jointly selected(van de Haar et al., 2019). To rule out this possibility, we carried out a reanalysis of conditional selection in cancer types with sufficient sample size and well-defined molecular subtypes, stratifying the data according to subtype classification. Although the reduction in sample size affected the ability to detect conditional pairs as statistically significant, we recovered some of the key interactions that characterized the modules of the epistasis network (Supplementary Table 4). More generally, we found that the estimated effect sizes for conditional selection in individual subtypes were highly correlated with the values obtained in the whole dataset (Figure 5a). Altogether, the results of the stratified analysis show that conditional selection also occurs within molecular subtypes and, therefore, is not an artifact of the joint analysis of samples from different groups.

**Figure 5:**
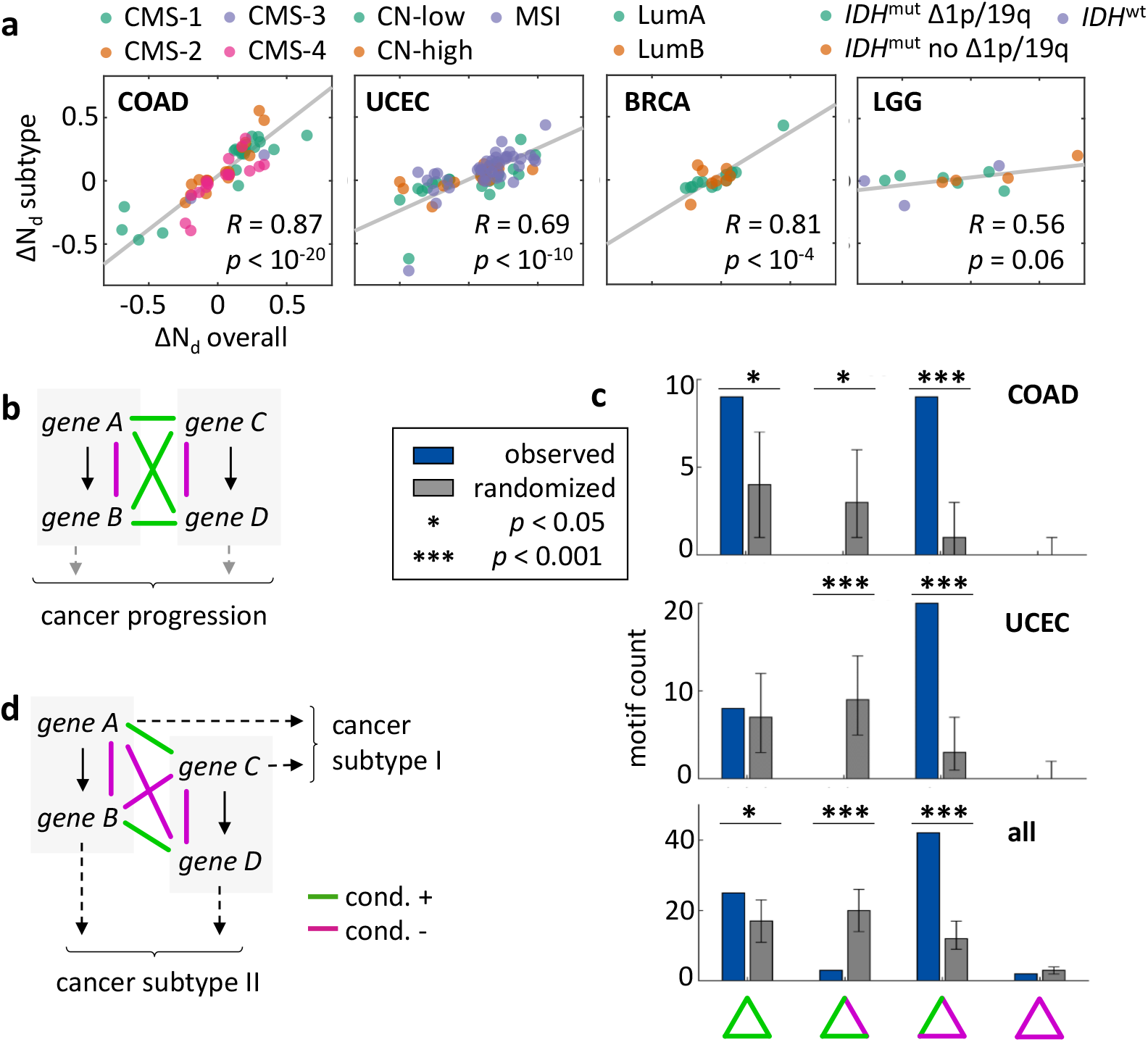
Conditional selection and structure of cancer molecular subtypes. a) Effect sizes of conditional selection at the cancer type (x axis) and subtype (y axis) levels. Each data point corresponds to a gene pair tested in at least one subtype. Gray lines represent the orthogonal least squares fit for each cancer type. b) Simple scenario of cancer progression through the alteration of two independent pathways (each represented in a gray rectangle), generating a pattern of within-pathway mutual exclusivity and across-pathway co-occurrence. c) Number of signed triangular motifs in the epistasis networks of colon cancer (COAD), endometrial cancer (UCEC), and all cancer types. Gray bars correspond to randomized networks with the same size and signed degree distributions obtained by pairwise rewiring of the network edges, and serve as a null model for motif counts (error bars represent 95% confidence intervals based on 2000 randomizations). BRCA: breast cancer; LGG: low-grade glioma. d) A more complex scenario of cancer progression, compatible with the overrepresentation of (+,-,-) triangular motifs. In this scenario, gene-specific, across-pathway synergistic interactions can lead to the development of distinct cancer subtypes.

Mutual exclusivity and co-occurrence among cancer somatic mutations have traditionally been interpreted in terms of within-pathway redundancy and across-pathway synergy(Yeang et al., 2008). In the simplest scenario, the synergy involves activation of an oncogenic pathway that promotes proliferation coupled with inactivation of a tumor suppressor pathway that regulates a cell cycle checkpoint(Cui et al., 2007). A relevant question concerning across-pathway synergy of driver mutations is whether it occurs in a combinatorial (or gene-agnostic) manner, that is, regardless of the specific genes that are mutated in each pathway(Oikonomou et al., 2014). To address this question, we investigated the distribution of triangular motifs in the epistasis networks. If across-pathway synergy was gene-agnostic, positive epistasis would manifest as an excess of triangular motifs containing two positive and one negative connection (Figure 5b). Contrary to this expectation, we found a significant excess of triangular motifs containing one positive and two negative connections in the epistasis networks (Figure 5c). Such configuration suggests a more complex scenario, in which across-pathway synergy is gene-specific, that is, depends on which genes are mutated in each pathway (Figure 5d). In that scenario, mutual exclusivity among functionally redundant mutations in the same pathway can still be expected; however, only some combinations of mutations affecting different pathways can act synergistically driving the progression of specific cancer subtypes. Consistent with that scenario, visual inspection of epistasis networks reveals a highly gene-specific pattern of across-pathway interconnections, whereby genes from the same pathway (for example, *RNF43* and *APC*, or *BRAF* and *RAS*) rarely display positive epistasis with the same partners. This finding calls for a reappraisal of the adequacy of pathway-centric approaches to cancer molecular biology. Although focusing on cancer pathways rather than individual genes can simplify the overall picture and boost the sensitivity of genomic analyses, it comes at the cost of overlooking the possibility that driver mutations in different genes from the same pathway can have distinct effects on tumor progression due to their involvement in different interactions with other pathways.

To better understand the effect of epistasis on cancer mutation landscapes, we simulated tumor evolution in the presence of conditional selection (Figure 6a). The simulation results suggest that the structure of epistasis networks can play a major role in determining the set of driver mutations observed in evolved tumors. In particular, highly modular facilitation networks with across-module interference, such as those observed in low grade gliomas, colorectal, and endometrial cancers, spontaneously give rise to clearly differentiated mutation-based subtypes (Figure 6b, column 4). This phenomenon is less pronounced in star-like epistasis networks (that is, those characterized by a single master driver that enhances the fitness of several secondary drivers; Figure 6b, columns 2-3) and completely absent in negative-modular networks (those corresponding to an idealized scenario of full within-pathway redundancy and across-pathway synergy; Figure 6b, column 5).

**Figure 6:**
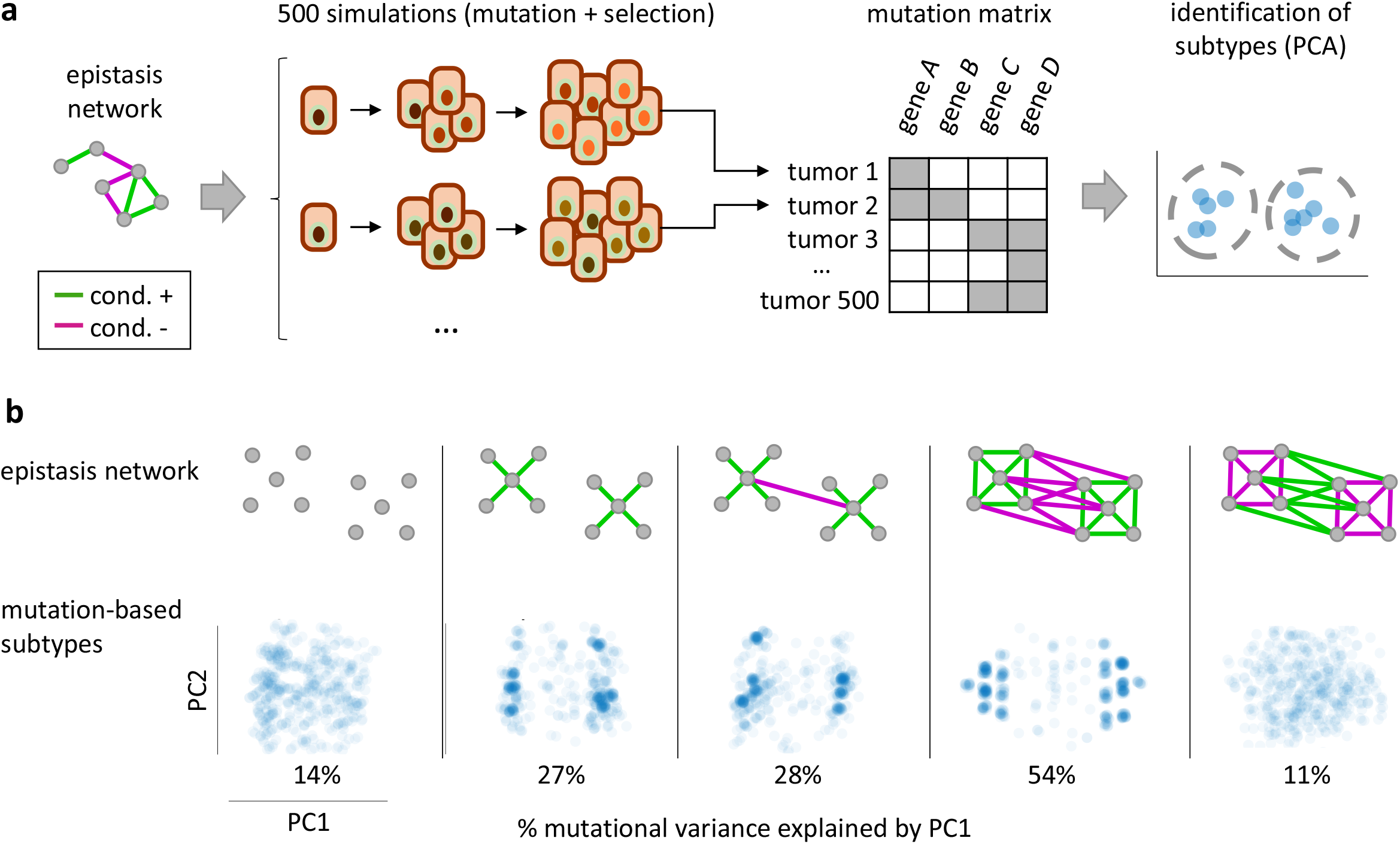
Modular epistasis in simulated networks leads to differentiation of cancer genomes into mutational subtypes. a) Tumor evolution was simulated in the presence of conditional selection, with epistatic interactions determined by networks with different degrees of modularity. For each network, the mutation composition of 500 simulated tumors was explored using principal component analysis to search for mutational differentiation into subtypes. b) Conditional selection can generate clearly delineated mutational subtypes if the underlying epistasis network (top) is characterized by intra-module facilitation and inter-module interference. The bottom panels show the results of a principal component analysis for each network, with each point representing a simulated tumor.

Based on these results, we hypothesize that tumor evolution is constrained by the structure of epistasis networks. In cancer types with strong modular epistasis, early drivers could be major determinants of the evolutionary path that a precancerous clone will take by modifying the strength of selection on subsequent mutations. Eventually, such evolutionary path can lead to an advanced tumor in which the mutational landscape is characterized by drivers in genes from the same epistasis module as the original one. Accordingly, differences in the pervasiveness of conditional selection could underlie the existence of clearly delineated (sub)type-specific gene sets in some cancer types, such as gliomas, colorectal, and endometrial cancers, but not in others, such as lung and liver cancers (Iranzo et al., 2018). The hypothesis that subtypes are shaped by conditional selection is consistent with previous findings(Zhang et al., 2017) and could open a new avenue for forecasting cancer evolution, encouraging further research as new somatic mutation datasets for precancer states and early cancers become available.

### Conditional selection affects survival rates

To assess the clinical relevance of conditional selection among cancer genes, we investigated the association between the modules of the epistasis network and patient survival times (Figure 7). To that end, we assigned tumor samples to epistatic modules by selecting, for each sample, the module that harbored the highest number of coding mutations. Module assignment was significantly associated with differential survival in 5 cancer types (glioblastoma, lower grade glioma, head and neck squamous carcinomas, pancreas adenocarcinoma, and endometrial cancer). An association between conditional selection and differential survival was also observed at the gene pair level, although the statistical significance was lower due to the small sample sizes (Suppl Figure 3). Of the 80 gene pairs with sufficient number of samples (at least 10 double mutants and 10 single mutants for each gene), we identified 9 instances (11% of the pairs), in which patient survival differed between tumors carrying coding mutations in both genes and those mutated in each gene alone (Cox regression and log-rank tests with p<0.1 for the omnibus test and p<0.15 for double-vs single-mutant comparisons). For comparison, we ran the same analysis on non-conditionally selected gene pairs, finding that only 4.6% (27 out of 580) showed differences in patient survival between double and single mutants. Compared to single-gene mutants, the survival of patients with double mutations can be improved, worsened, or unaffected. Notably, worse patient survival is associated with gene pairs subject to positive conditional selection (*ARID1A-ERBB3* in bladder cancer, *CASP8-FAT1* in head and neck cancers, *FGFR2-TP53* in endometrial cancer, and *MUC17-NF1* in melanoma), whereas improved patient survival is associated with negative conditional selection (*PPP2R1A-PTEN* and *ARID1A-PPP2R1A* in endometrial cancer). The association with better patient survival in *PPP2R1A-PTEN* and *ARID1A-PPP2R1A* double mutants holds when controlling for mutational load, and suggests that mutual exclusivity in these gene pairs could result from synthetic lethality. Likewise, synthetic lethality might underlie other instances of negative conditional selection that could not be tested due to a lack of double mutants, as has been shown for *KRAS* and *EGFR* in lung cancer(Unni et al., 2015).

**Figure 7:**
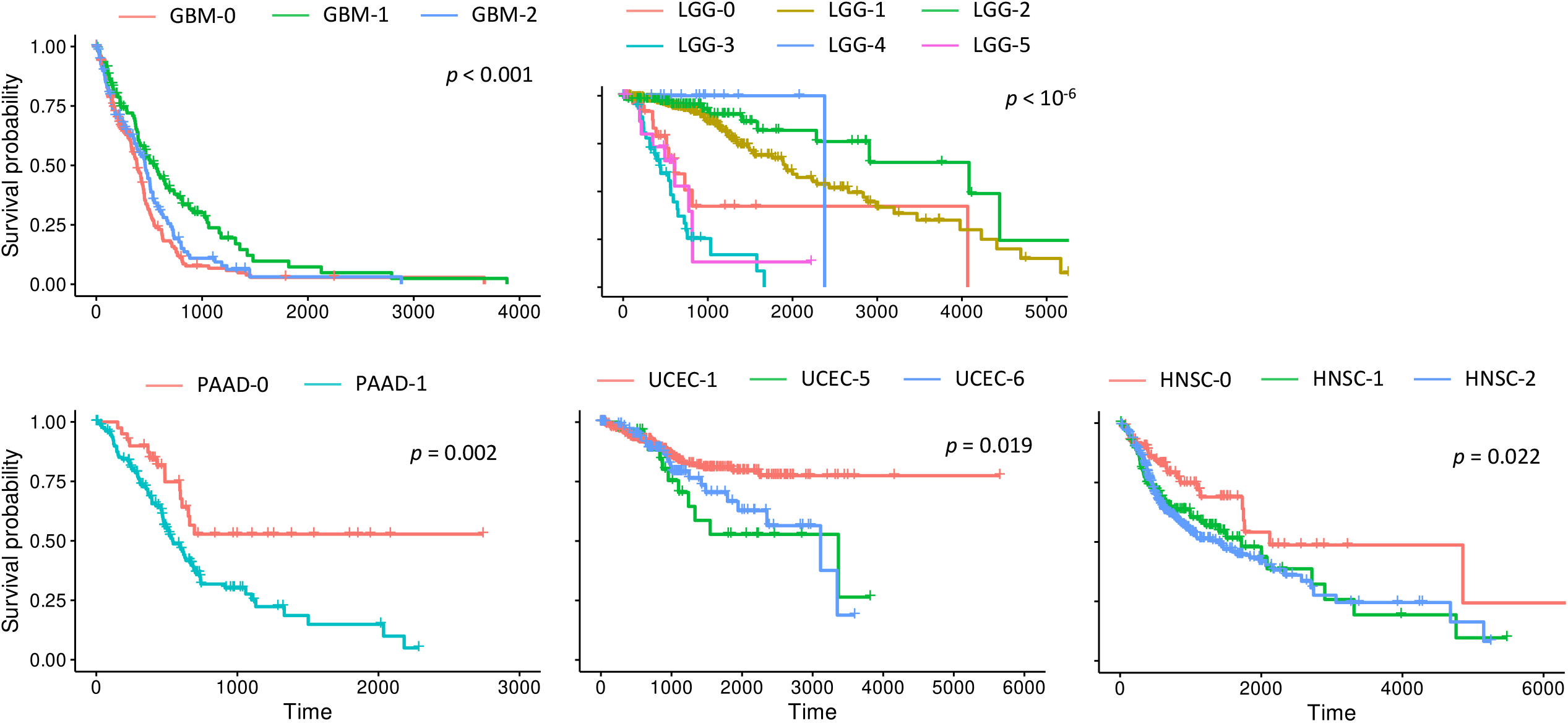
Differences in patient survival associated to the modules of the epistasis network. Patients were assigned to the module with which they shared the greatest number of mutated genes. Patients without mutations in genes from the epistasis network were assigned to a null module (e.g., GBM-0). P-values correspond to the log-rank test for the null hypothesis that there are no differences among the three curves. GBM: glioblastoma; LGG: lower grade glioma; PAAD: pancreatic cancer; UCEC: endometrial cancer; HNSC: head and neck squamous carcinoma. Genes included in each module GBM-1: *ATRX, IDH1, RB1, TP53*; GBM-2: *PTEN*; LGG-1: *ATRX, IDH1, TP53*; LGG-2: *CIC, FUBP1*; LGG-3: *EGFR, PTEN*; LGG-4: *IDH2*; LGG-5: *NF1*; PAAD-1: *KRAS, TP53*; UCEC-1: *ARID1A, CTCF, CTNNB1, INPPL1, JAK1, KANSL1, KMT2B, KMT2D, KRAS, NFE2L2, PIK3R1, PTEN, RB1, RNF43, TAF1, ZFHX3, ATM, CCND1, FGFR2, UPF3A*; UCEC-5: *PIK3CA*; UCEC-6: *PPP2R1A, TP53*; HNSC-1: *CASP8, CTCF, EP300, EPHA2, FAT1, FBXW7, HLA-A, HLA-B, HRAS, RAC1, TGFBR2*; HNSC-2: *CDKN2A*.p14arf, *CDKN2A*.p16INK4a, *TP53*.

These results are of particular interest in the context of personalized medicine, as they show that epistasis-driven mutational profiles could be better predictors of cancer prognosis than single mutations. Conceivably, epistasis profiles, such as those described here, could be leveraged to aid the selection of optimal therapeutic strategies. In particular, the set of epistatic interactions, in which a driver mutation is involved, could be indicative of the mechanism, through which that driver contributes to the fitness of the tumor. Thus, epistasis profiles could be key for understanding the variability in the effect of gene-targeting therapies across patients and cancer subtypes.

### Limitations of the study

An obvious limitation of the approach developed in this work is that we only sought to identify pairwise interactions between driver mutations. Extending *Coselens* to search for higher-order interactions is straightforward. However, because quantifying conditional selection while accounting for heterogeneity in mutational mechanisms requires partitioning the data in as many groups as conditions, and the number of conditions grows exponentially with the interaction order, the detailed study of high-order interactions can only be performed in large datasets and for frequently mutated cancer genes.

Another limitation is that we only analyzed point mutations that lead to amino acid replacement. Extending the approach to other types of mutations, in particular Copy Number Alterations (CNA), will require developing the corresponding mutational models that, in the case of CNA, will have to take into account correlations among adjacent genes. We anticipate that, by including CNA in future studies, additional instances of conditional selection will be identified for tumor suppressor genes that are typically affected by homozygous deletions (Ciriello et al., 2012; Mina et al., 2017). It also seems plausible that positive selection for CNA in tumors with chromosome instability is associated with inactivating mutations in the p53 pathway given that the latter facilitates the survival of cells with genomic aberrations (Hanel and Moll, 2012; Motoyama and Naka, 2004).

A third aspect that deserves further investigation concerns the temporal dynamics of epistasis. Changes in the mutation landscape and the selective pressures experienced by the tumor likely modify the structure of epistasis networks in different stages of cancer progression (Persi et al., 2021). However, due to the transversal nature of the TCGA, the dynamic nature of epistasis was not captured in this study. Future analyses of somatic mutations in healthy tissues and precancerous lesions would be of great interest, as they could lead to the discovery of new epistasis modules associated with protection against cancer progression.

## Conclusions

In this work, we developed a method, Coselens (COnditional SELection on the Excess of Nonsynonymous Substitutions), for analysis of epistasis in cancers. Coselens includes two major features that, to the best of our knowledge, are missing in methods previously developed for the same purpose, namely, rigorous control for biases introduced by cancer subtypes and explicit estimation of the size of epistatic effects. The results reported here expand the catalogue of conditionally selected pairs of cancer genes by several folds and demonstrate wide spread and importance of conditional selection in cancer evolution. Indeed, in some cancer types, up to 50% of driver mutations are involved in such epistatic interactions. Furthermore, we observed association between conditional selection and patient survival, attesting to the clinical relevance of epistasis. Perhaps, most notably, the findings of this work emphasize the striking complexity of the epistasis among driver mutations that is manifest at several levels. First, we identified multiple cases of all theoretically conceivable forms of epistasis, both positive and negative, including strict dependence, facilitation, interference, and masking. Second, many of the conditionally selected gene pairs are cancer subtype-specific and organize into highly modular epistasis networks. Third, we found that synergistic epistasis between signaling pathways is highly gene-specific. Conceivably, epistasis shapes differentiated cancer subtypes by constraining the evolutionary path that follows early driver mutations. This evolutionary path towards advanced tumors involves mutations in a specific set of additional genes that belong to the same module of the epistasis network and determine the key properties of a molecular cancer subtype. Finally, the finding that epistatic interactions involving widespread cancer genes are cancer type-dependent raises the possibility that the same gene might act through different mechanisms in different cancer types. Thus, epistasis profiles potentially could guide the application of gene-targeting therapies across cancer types.

## Acknowledgements

We thank Iñigo Martincorena and Koonin group members for helpful discussions. This work was initiated during an internship of G.G. at Koonin group, which was made possible thanks to the NIH Summer Internship Program in Biomedical Research. This work utilized the computational resources of the NIH HPC Biowulf cluster (http://hpc.nih.gov).

J.I. is supported by the Ramón y Cajal Programme of the Spanish Ministry of Science (Grant No. RYC-2017-22524), the Agencia Estatal de Investigación of Spain (Grant No. PID2019-106618GA-I00), and the Severo Ochoa Programme for Centres of Excellence in R&D of the Agencia Estatal de Investigación of Spain (Grant No. SEV-2016-0672 (2017–2021) to the CBGP). G.G. is supported by the DHHS/PHS/National Institutes of Health (Grant No. 2R01DE019637-10, “Patterning the vertebrate dentition through replacement and repair). J.C.-E. is supported by the Youth Employment Initiative of the European Social Fund through a junior postdoctoral contract from Comunidad de Madrid (Grant No. PEJD-2019-POST/BIO-16377). E.V.K. is supported by intramural research program funds of the National Institutes of Health (National Library of Medicine).

## Author contributions

J.I. and E.V.K. designed and supervised the study; J.I. and G.G. developed the *coselens* software and generated the primary results; J.I., G.G., and E.V.K. analyzed the results; J.C.-E. developed and simulated the computational model of tumor evolution; J.I., G.G., J.C-E., and E.V.K. wrote the paper.

## STAR Methods

### Dataset and data preprocessing

Somatic mutation calls for The Cancer Genome Atlas (TCGA) dataset, generated by the Multi-Center Mutation Calling in Multiple Cancers project(Ellrott et al., 2018), were downloaded from the Genomic Data Commons repository of the National Institutes of Health (https://gdc.cancer.gov/about-data/publications/mc3-2017). Clinical and survival data for TCGA were also downloaded from the Genomic Data Commons repository of the National Institutes of Health (https://gdc.cancer.gov/node/905/; file TCGA-CDR-SupplementalTableS1.xlsx)(Liu et al., 2018). To avoid the confounding effect of purifying selection, which is non-negligible in hypermutated tumors(Persi et al., 2018), samples with >3000 coding mutations were classified as hypermutators and removed from subsequent analyses.

### Inference of conditional selection: the Coselens method

To quantify conditional selection at the gene level, we developed *Coselens* (*COnditional SELection on the Excess of Nonsynonymous Substitutions*), a tool that makes extensive use of the *dndscv* R package (dndSCV, RRID:SCR_017093) to quantify selection in cancer and normal tissues(Martincorena et al., 2017). We briefly describe the fundamentals of the *dndscv* method, as long as it is needed to understand the modifications introduced by *Coselens*; we refer the reader to the original publication for further details. In short, *dndscv* estimates the number of missense and truncating (nonsense and essential splice site) substitutions per gene that would be expected in the absence of selection by combining a nucleotide substitution model with 192 rates (12 possible substitutions in each of the 16 possible trinucleotide contexts) and a set of genomic covariates, which greatly improve the performance of the method at low mutation loads. The model parameters are estimated through negative binomial regression on the number of synonymous mutations to maximize the following likelihood function:

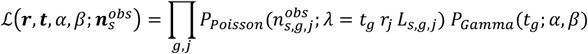

where 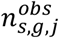 is the number of synonymous substitutions observed in the gene *g* and the index *j* corresponds to the substitution classes (192 by default although substitution models with fewer or more classes can be easily accommodated), *r*_*j*_ is the mutation rate for each substitution class, *L*_*s,g,j*_ is the number of sites in gene *g* that are synonymous under the substitution class *j*, and *t*_*g*_ is a factor accounting for variable mutation rates across genes, which is assumed to follow a gamma distribution with parameters *α* and *β*. Once those parameters have been inferred, the expected number of nonsynonymous substitutions in the absence of selection is obtained by replacing *L*_*s,g,j*_ by the number of sites in gene *g* that are susceptible to missense and truncating substitutions (*L*_*m,g,j*_ and *L*_*t,g,j*_, respectively). The selection parameter *ω*_*k,g*_ is then calculated for each gene as the ratio between the observed number of mutations and their neutral expectation 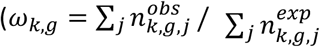, where the subscript *k* ∈ {*m, t*} refers to missense and truncating substitutions). From the perspective of the likelihood function, the selection parameter is a multiplicative factor in *λ* that modifies the observed substitution frequency with respect to the neutral expectation, with *ω*_*k,g*_ > 1 and *ω*_*k,g*_ < 1 implying positive and purifying selection, respectively. The significance of *ω*_*k,g*_ is computed through a likelihood ratio test, where the null and alternative hypotheses correspond to *ω*_*k,g*_ = 1 and *ω*_*k,g*_ ≠ 1, respectively.

To develop *Coselens*, we modified the *dndscv* method in two substantial ways. First, we quantified the differences (rather than the ratios) between the observed number of missense and truncating substitutions and their neutral expectations:

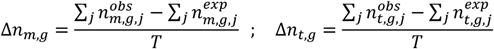

where *T* is the number of samples used in the analysis. Missense and truncating substitution excesses result from mutations that have reached detectable frequencies through positive selection. Because the effect of purifying selection in non-hypermutator tumors is negligible(Martincorena et al., 2017; Persi et al., 2018), the total excess of nonsynonymous mutations in a gene, *Δn*_*g*_ = Δ*n*_*m,g*_ + Δ*n*_*t,g*_, is a good estimate of the number of driver substitutions *N*_*d*_ in that gene (formally, 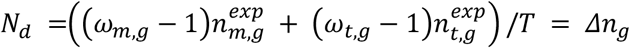; note that we omitted the subscript *g* in *N*_*d*_ for simplicity although it is a gene-specific quantity). Second, we modified the likelihood ratio test to allow for comparison of *Δn*_*g*_ between two sets of samples. To this end, *Coselens* separately fits the *dndscv* substitution model to each sample group (hereafter *X* and *Y*), obtaining two sets of parameters for the neutral scenario (***θ***(*X*) and ***θ***(*Y*)) and two sets of nonsynonymous mutation excesses (Δ*n*_*k,g*_(*X*) and Δ*n*_*k,g*_(*Y*), with *k* ∈ {*m, t*}). The likelihood ratio test is built according to the following null and alternative hypotheses (2 degrees of freedom):

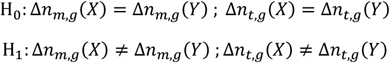

To restrict the hypothesis test to differences in the number of drivers, but not in the underlying mutation rates, the null hypothesis does not introduce any constraint on the parameters ***θ***(*X*) and ***θ***(*Y*). Accordingly, the likelihood of the null hypothesis is evaluated using group-specific parameters for each observation and joint nonsynonymous mutation excesses Δ*n*_*m,g*_(*X* + *Y*) and Δ*n*_*t,g*_(*X* + *Y*)), for which the general expression is

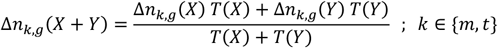

Here, *T*(*X*) and *T*(*Y*) are the numbers of samples in each group. The joint nonsynonymous excesses can be incorporated in the likelihood function by leveraging the relationship between the mutation excess Δ*n*_*k,g*_ and the mutation ratio *ω*_*k,g*_, that is,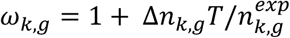. Under the null hypothesis, this relationship becomes

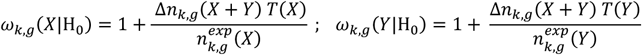

Thus, the likelihood of the null hypothesis becomes

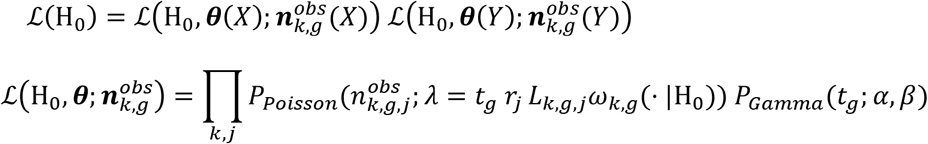

The expression for the likelihood of the alternative hypothesis is analogous, except that the *ω*_*k,g*_ parameters are calculated using sample-specific mutation excesses, that is,

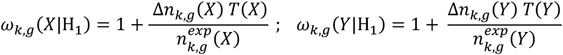

The output of the likelihood ratio test is the probability that the excess of nonsynonymous substitutions (missense, truncating, or both) in a given gene differs between the two sets of samples. As mentioned above, in the context of nearly absent purifying selection, such as in cancer evolution, the effect size can be interpreted as the difference in the number of driver mutations per sample in the gene of interest:

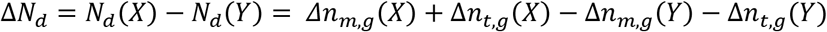

In our case, the sets *X* and *Y* correspond to samples with and without coding mutations in a second gene; however, the *Coselens* method is general and can be applied to compare any pair of sample sets (e.g. to study differential selection across tissues or clinical conditions). Although we focused on genes under positive selection, *Coselens* can also detect differences in purifying selection if the excess of substitutions is negative. To achieve the highest accuracy, *Coselens* should be run on whole exome data or targeted sequencing data including as many genes as possible, as that boosts the performance of parameter inference. The output can be subsequently filtered to restrict false discovery rate correction and downstream analysis to a set of genes of interest.

We provide *Coselens* as an open-source R package that can be downloaded from https://github.com/ggruenhagen3/coselens. Besides the main method described above, the package includes some extensions to study the excess of small indels and test for differences in missense or truncating substitutions only.

### Conditional selection within cancer types

To investigate pairwise conditional selection among cancer genes, we compiled a set of candidate gene pairs, to which we would apply the *Coselens* method. Each candidate gene pair consisted of a *split gene* (the gene inducing conditional selection) and a *query gene* (the gene that experiences conditional selection). Candidate gene pairs were separately identified in each cancer type; in that sense, one can think of gene pairs as triplets (cancer type – split gene – query gene). Given a split gene, conditional selection was assessed by dividing the samples into those that carried coding mutations (nonsynonymous substitutions and small indels) in the split gene and those that do not, comparing both groups with *Coselens*, and filtering the results according to the list of query genes associated to that split gene.

A preliminary set of query genes was built by finding genes evolving under significant positive selection in each cancer type with the R package *dndscv* (default parameters). From this preliminary set, we built the set of split genes that harbored coding mutations in at least 25 samples, while being free of coding mutations in at least another 25 samples. Then, we expanded the set of query genes by running *Coselens* and searching for conditional selection on known cancer genes not included in the preliminary set of query genes. For this exploratory step, we considered a list of 369 cancer genes from previous studies(Iranzo et al., 2018; Martincorena et al., 2017) and applied the Benjamini-Hochberg false discovery rate correction to all possible pairs consisting of a split gene and one of those known cancer genes. This approach identified 20 additional query genes, for which positive selection is only detectable in a restricted subset of samples (e.g., *HLA-C* and *PGM5* in *BRAF*-mutant colorectal cancer). We finally studied conditional selection on all possible split - query gene pairs within each cancer type, applying a false discovery rate correction to the pooled results (that is, jointly considering all cancer types). Additionally, to capture cases of reciprocal conditionality among gene pairs that could have been missed due to insufficient statistical power, we applied restricted hypothesis testing to split-query gene combinations whose reciprocal had been found significant under the standard approach.

### Negative control

A control dataset was built by randomly assigning mock “mutated split gene” and “wild-type split gene” labels to each tumor. Relabeling was done such that the total number of mutations in the control subsets remained approximately the same as in the original ones. Specifically, for each cancer type, tumor samples were randomly chosen and assigned to the “mutated split gene” subset until the total number of mutations in the group reached or surpassed the number of mutations in the original subset of samples with mutations in the split gene. The reason to keep the number of mutations (rather than the number of samples) per group fixed is that *Coselens* pools together all mutations from all samples within a group. By keeping that number approximately unchanged, we ensured that the sensitivity of *Coselens* was similar in the negative control and in the original dataset. Furthermore, it is important to perform the randomization at the sample (rather than mutation) level to conserve the structure of within-sample correlations, so that the model covariates remain informative. Note that, because this randomization procedure removes systematic biases in mutational load and subtype composition between subsets, it is not suitable to assess the rate of false positives induced by such confounding factors. The entire workflow described in the previous section was repeated for 40 randomized control datasets, finding an average of 6.5 significant pairs per run (0.1% of the tested pairs). The comparison between this number and the value found in the original dataset (296 significant pairs) yields an empirical false discovery rate of ∼2%.

### Conditional selection within cancer subtypes

Samples from colorectal cancer were classified into molecular subtypes, and each subtype was separately analyzed as described above. The consensus molecular subtypes of colorectal cancer samples were obtained from the data repository of the colorectal cancer subtyping consortium (https://www.synapse.org/#!Synapse:syn2623706/files/)(Guinney et al., 2015). After removing hypermutated tumors (>3000 coding mutations), only 49 samples remained in the CMS1 group. Therefore, the sample size threshold to consider a candidate gene as a split gene in CMS1 was lowered to at least 18 samples with and 18 samples without coding mutations in that gene. A similar subtype-based analysis was applied to endometrial cancer, low-grade glioma, and breast cancer. The molecular subtype classification for those samples was obtained from the original publications(Cancer Genome Atlas Research et al., 2013; Ceccarelli et al., 2016) and the PanCancer Atlas dataset(Berger et al., 2018) of the cBioPortal repository(Cerami et al., 2012).

### Classification of conditional pairs by into modes of epistasis

Gene pairs were classified by epistasis modes attending to the variation in the excess of nonsynonymous mutations in the first gene (*A*) when the second gene (*B*) does or does not harbor coding mutations (*N*_*d*_(*A*|*B*) and 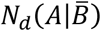, respectively). First, we tested whether *N*_*d*_ (*A*|*B*) or 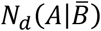were significantly distinct from zero. Within the *dndscv* framework, testing *N*_*d*_ ≠ 0 is equivalent to testing whether the ratio *dn*/*ds* ≠ 0. To account for multiple comparisons, we applied the Benjamini-Hochberg correction to all pairs tested for conditional selection and set the significance threshold to a false discovery rate of 5%. Next, we classified conditionally selected gene pairs in one of the following categories: (i) masking epistasis, if 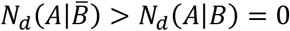; (ii) interference, if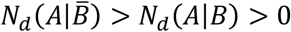; (iii) facilitation, if 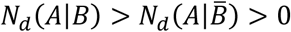; (iv) strict dependence, if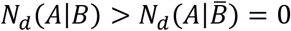; (v) strict dependence with sign change, if 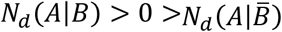; (vi) compensation, if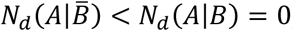; (vii) conditional deleterious, if 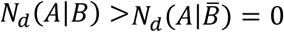; and (viii) incompatible, if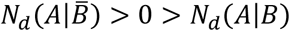. Because the test for conditional selection can be more sensitive than the test for *N*_*d*_ ≠ 0, there were some cases where 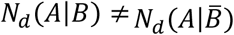 but neither *N*_*d*_(*A*|*B*) nor 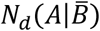were significantly different from zero. Those cases were left unclassified.

### Estimation of the number of driver substitutions affected by conditional selection

For a given cancer type and a split gene *B*, the average number of driver events per tumor that involve conditional selection on the query gene *A* was obtained as Δ*N*_*d*_(*A, B*) *X*_*B*_, where 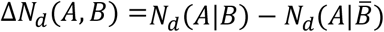is the effect size of conditional selection and *X*_*B*_ is the fraction of tumors with coding mutations in the split gene *B*. To estimate the total number of driver events affected by conditional selection, we summed over all directed pairs (*A, B*) that were significant with FDR ≤ 5%, including reciprocal pairs that were significant under restricted hypothesis testing.

### Estimation of the number of driver mutations per patient

Although the number of driver mutations can be estimated within the *dndscv* (and *Coselens*) framework, we chose an independent approach to minimize the risk that correlations with the number of conditional driver substitutions were affected by the presence of uncontrolled confounding factors. Therefore, we implemented a previously described simple regression-based approach(Iranzo et al., 2018). The number of driver mutations per patient was estimated as the intercept of a linear regression model that had, as independent variables, the cancer type, the number of coding mutations in passenger genes, and the interaction of both; and as the dependent variable, the number of coding mutations in 441 significantly mutated cancer genes (the union set of those previously detected(Martincorena et al., 2017) and those found in this study).

### Functional association between genes in conditionally selected pairs

Each split-query gene pair was searched against the STRING database of protein interactions (v11.0). If an interaction was found in the database, the pair was assigned a score *s* equal to the STRING score divided by 1000 (this normalization accounted for the fact that STRING scores range from 0 to 1000, with higher values representing higher confidence in the predicted interaction). Pairs not found in the STRING database were assigned a score of zero. Given a subset *G* of gene pairs, the STRING interaction ratio was calculated as *r*_*G*_ = ∑_*G*_ *s* / ∑_*G*_(1 − *s*). In the special case when *s* takes a binary value (1 or 0), *r*_*G*_ becomes the ratio between the number of gene pairs that are known to interact and those for which there is no evidence of interaction. The STRING interaction ratio was computed for increasing magnitudes of conditional selection by defining a nested collection of subsets *G*_*x*_, with 0 < *x* < max(Δ*N*_*d*_), such that each subset includes all gene pairs whose effect size Δ*N*_*d*_ is greater than *x* (accordingly, *G*_0_ corresponds to the set of all gene pairs included in the analysis). The interaction enrichment ratio for a given effect size was then calculated as *R*(Δ*N*_*d*_) = *r*_*G* Δ*N*_*d*/*r*_*G* 0_. Conceptually, the enrichment ratio is analogous to a risk ratio, with the difference that the response variable is continuous rather than binary. Following Pagano and Gauvreau (2000), the 95% confidence interval was calculated as *CI*_95_(Δ*N*_*d*_) = exp (ln(*R*) ± 1.96 × *SE*(ln(*R*))), where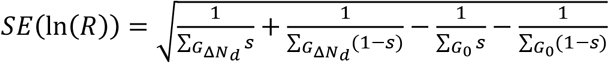.

### Reciprocity of conditional selection

Let 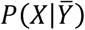be the conditional probability that query gene *X* harbors a driver mutation provided that there are no driver mutations in the split gene *Y*. Conversely, let *P*(*X*|*Y*) be the conditional probability that the query gene *X* harbors a driver mutation provided that there are driver mutations in the split gene *Y*. Then, the magnitude of epistasis-driven conditional selection can be quantified as the difference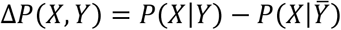. For any pair of genes *A* and *B*, we are interested in finding an expression that relates Δ*P*(*A, B*) and Δ*P*(*B, A*), that is, the magnitude of conditional selection when *A* is the query gene and *B* is the split gene, and when the query and split roles are reversed. Combining Bayes theorem and the identity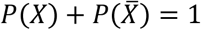, the difference Δ*P*(*A, B*) can be expressed as

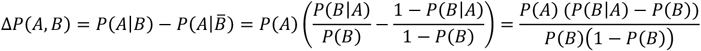

where *P*(*A*) and *P*(*B*) are the marginal probabilities for finding driver mutations in genes *A* and *B*, respectively. In a similar way, Δ*P*(*B, A*) can be written as

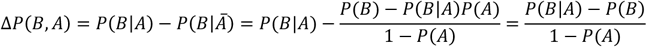

Extracting *P*(*B*|*A*) − *P*(*B*) from both expressions, and after some algebraic manipulation, we obtain

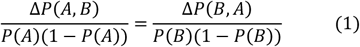

If presence or absence of driver mutations in genes *A* and *B* are modeled as independent Bernoulli random variables with success probabilities *P*(*A*) and *P*(*B*), the denominators in eq. (1) correspond to the variances of such random variables across tumors. Accordingly, the left and right terms of eq. (1) can be interpreted as the standardized effect sizes of conditional selection for the query genes *A* and *B*, respectively. Eq (1) implies that, if conditional selection results from epistasis between driver mutations, the standardized effect sizes should be equal for reciprocal query-split gene pairs.

Qualitatively, reciprocity implies that, if conditional selection is detected for a given split-query gene pair, it should be also detected for the reverse pair, in which the split and query genes are exchanged. Attending to this qualitative criterion, we found that 68% of the pairs that are subject to significant conditional selection are reciprocal (this fraction does not include reverse pairs that could not be tested due to insufficient number of coding mutations in one of the genes).

To carry out a more quantitative comparison between the theoretical prediction with the empirical estimates, we first established a correspondence between the marginal and conditional probabilities from eq. (1) and the nonsynonymous mutation excesses calculated from cancer genomic data. To that end, we assumed that only the first driver substitution in each cancer gene is positively selected, and therefore the probability to find a driver mutation in a given gene can be approximated by the mean number of driver substitutions per tumor in that gene. Although this assumption is strictly valid only for single-hit oncogenes, it also holds for tumor suppressor genes, as long as the inactivation of the second copy of the gene can occur via large deletions. Thus, to validate eq. (1) with empirical genomic data, we replaced Δ*P*(*A, B*) ≈ Δ*N*_*d*_(*A, B*) and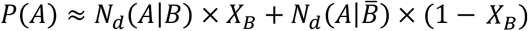, where *X*_*B*_ is the fraction of tumors with coding mutations in the split gene. This second approximation potentially could become a major source of error because the empirical split of tumor samples is based on the presence of coding mutations, whereas conditionality in the theoretical expression specifically involves driver mutations. Therefore, calculating the goodness of fit of the empirical data to the theoretical expectation in eq. (1) is a straightforward way to assess whether the empirical estimates are robust with respect to the use of coding mutations rather than driver mutations in the split gene to quantify conditional selection.

### Assessment of the effect of differential mutation availability on conditional selection estimates

To investigate whether conditional selection can be explained by differences in mutation availability, two alternative approaches were adopted. The first approach consisted of inferring, given the expected number of mutations per tumor in the query gene, the probability that at least one driver mutation becomes available for selection to act on. We considered a Poisson distribution for the number of substitutions per gene per tumor, such that given a local mutation rate μ, the probability that at least one driver mutation becomes available in the gene of interest is *π*(*µ*) = 1 − exp(*bµ*). Given a set of *T* independent tumors, the number of tumors in which the driver mutation becomes available (*k*) follows a binomial distribution with parameters *T* and *π*(*µ*). Following the central limit theorem, for large enough *T*, the fraction of tumors in which the driver mutation becomes available (*f* = *k*/*T*), can be approximated by a normal distribution with mean *π*(*µ*) and variance *π*(*µ*)(1 − *π*(*µ*))/*T*. Therefore, given two sets of values for *f, µ*, and *T* (corresponding to tumors with and without mutations in the split gene), it is possible to apply a maximum likelihood approach to infer the only unknown parameter of the model (*b*) and then obtain the probability that differences in *f* can be explained by changes in μ. To perform these calculations, we took the expected number of non-synonymous and non-sense mutations per sample per gene inferred with the dndscv method as a gene-wise measure of μ; and the average number of drivers per gene per tumor as a gene-wise measure of *f* (i.e., *f* ≈ *N*_*d*_, following the same rationale described above for the analysis of reciprocity). A continuity correction was included when assessing the likelihood, such that *L*(*b*; *f*, μ, *T*) = NormCDF(*f* + 0.5/*T*) – NormCDF(*f -* 0.5/*T*), where “NormCDF” is the cumulative probability function for a normal distribution with mean *π*(*µ*) and variance *π*(*µ*)(1 − *π*(*µ*))/*T*. For each conditional gene pair, the probability that it can be explained by differential mutation availability was calculated as the product *p* = *p*_1_*p*_2_, where *p*_1_(*f*_1_, *µ*_1_, *T*_1_) is the probability that *f*_1_ (the average number of drivers per tumor in samples with a mutated split gene) comes from a normal distribution with mean *π*(*µ*_1_) and variance *π*(*µ*_1_)(1 − *π*(*µ*_1_))/*T*_1_ and *p*_2_(*f*_2_, *µ*_2_, *T*_2_) represents the same probability for samples without coding mutations in the split gene. In the case of reciprocal interactions, we combined the probabilities of the direct and reverse pairs (*p*^*AB*^ and *p*^*BA*^, respectively) as *p* = *p*^*AB*^*p*^*BA*^. The resulting probabilities were corrected for multiple comparisons with FDR = 5%.

The second approach involved a regression analysis to assess how the excess of gene pairs displaying positive epistasis (compared to those that display negative epistasis) changes with the difference in the median mutational load of tumors with and without mutations in the split gene. By focusing on the difference between positive and negative epistatic pairs, rather than on the total numbers, we accounted for the fact that mutational load has a directed effect on mutation availability, that is, higher (respectively lower) mutational loads increase (decrease) mutation availability, resulting in an apparent excess of positive (negative) epistatic pairs. We found a weak, but significant association between the difference in mutational load and the excess of positive epistatic pairs in which a split gene is involved (0.16 pairs per every 100 mutations/exome difference in samples with and without coding mutations in the split gene; ANCOVA stratified by cancer types, R^2^ = 0.15; p = 1e-9). To assess how this association could translate into apparent (mutation availability-driven) conditional selection, we adopted a Bayesian approach to infer the probability that a pair involving a given split gene can be attributed to differences in mutational load. First, we modeled the prior for the number of such pairs as a Poisson distribution with mean *λ* = *a* × Δ*µ*_*g*_, where *a* is the slope of the regression model and Δ*µ*_*g*_ is the difference in the median mutational load of tumors that do and do not harbor mutations in the split gene of interest. To obtain the posterior distribution, we took into account the total number of conditional interactions observed for that split gene, *k*_*g*_, by truncating the Poisson distribution between 0 and *k*_*g*_. The expected contribution of differential mutation availability to the number of conditional pairs was then calculated as the mean of the truncated distribution.

### Detection of modules in the epistasis networks

Positive conditional modules in the epistasis networks were identified with SiMap, a community detection algorithm that takes into account the sign and weight of interactions(Esmailian and Jalili, 2015). We used the effect sizes Δ*N*_*d*_(*A, B*) as interaction weights and preprocessed the networks to extract their connected components before running the community detection algorithm with default parameters. Overlapping modules were identified by adding, for each gene, the weights of its interactions with genes from other communities. A secondary module assignation was made for those genes for which the total weight of interactions with the original community was <1.5 times the total weight of interactions with the secondary module. Samples were assigned to the module with which they shared the greatest number of mutated genes. Samples without mutations in genes from the epistasis network were assigned to a null module. Ambiguous cases (samples with the same number of genes mutated in >1 module) were resolved only if they met the following two conditions: (i) there were only 2 candidate modules, to which the sample could be assigned, and (ii) the fraction of mutated genes belonging to each module was different. If both conditions were met, the sample was assigned to the module with the largest fraction of mutated genes; otherwise, the module assignation was considered ambiguous.

### Survival analysis

We conducted a survival analysis to investigate if the combination of mutations in two conditionally selected genes affected patient survival, compared to mutations in a single gene. For each gene pair (*A, B*) in a given cancer type, tumor samples were split in 3 groups: (i) those with coding mutations in gene *A* but not in gene *B*, (ii) those with coding mutations in gene *B* but not in *A*, and (iii) those with coding mutations in both genes (samples without mutations in *A* or *B* were not included). Gene pairs that did not reach a minimum of 10 samples in each group and 100 samples overall were no further considered. For those gene pairs that reached sufficient sample size, we adopted two approaches to assess the statistical significance of differences in survival between the double-mutation group and each of the single-mutation groups: (a) Kaplan-Meier estimation followed by non-parametrical log-rank tests(Clark et al., 2003), and (b) Cox regression analysis with the double-mutation group as the reference category(Bradburn et al., 2003). In both cases, we assessed the overall significance (omnibus test for the null hypothesis that survival is the same in the three groups) and the significance of the differences between each single-gene mutant and the double mutant. We reported the cases with p<0.1 for the omnibus test and p<0.15 for both single vs double mutant comparisons. Statistical analyses were carried out with the R packages *survminer* and *survival*(Therneau and Grambsch, 2000).

The association between patient survival and positive epistatic modules with >15 samples was assessed using the same approaches, and the cancer types reported in the text are those with p<0.05 for the omnibus tests.

### Comparison of Coselens and SELECT methods

The list of conditionally selected events in the TGCA identified using the *SELECT* algorithm was collected from the Supplementary Information of the original publication(Mina et al., 2017). To reconcile the fact that *SELECT* operates at the level of “selected functional events” (SFE) and *Coselens* operates at the gene level, each SFE was mapped to the gene or genes affected by that SFE. Pairs of conditional SFE that mapped to the same gene (typically associated with double inactivation of tumor suppressor genes) were not considered. Directionality was ignored and the significance threshold of both methods was adjusted to ensure a FDR <10%, which was the value originally used by the developers of *SELECT* when applying it to the TCGA dataset.

### Simulation of tumor evolution

The evolution of a population of (pre)cancer cells was modeled as a discrete time birth-and-death process, where death also accounted for differentiation into non-proliferative lineages. At each time step, each cell could either die (with probability *P*_*d*_) or divide (with probability 1 − *P*_*d*_). Following previous work by McFarland et al(McFarland et al., 2013), the death probability was set to

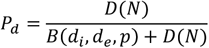

where *D*(*N*) = *N*/*K* represents density-dependent death rate (*N* is the current population size and parameter *K* determines the equilibrium population size in the absence of mutations). The birth term was modeled as

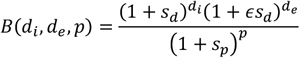

where *s*_*p*_ is the fitness effect of a passenger mutation, *s*_*d*_ is the fitness effect of a driver mutation in the absence of epistasis, and *ϵ* is an epistasis coefficient that accounts for the increase in the fitness effect of a driver mutation due to positive epistasis. The variable *p* represents the number of passenger mutations in the genome of a given cell, *d*_*i*_ is the number of driver mutations that are not affected by epistasis, and *d*_*e*_ is the number of driver mutations affected by positive epistasis. We assumed a scenario, in which negative epistasis fully masks the effect of driver mutations, and therefore, driver mutations affected by negative epistasis do not contribute to the birth term.

Upon division, one of the daughter cells can acquire driver and passenger mutations. The number of new driver and passenger mutations was drawn from Poisson distributions with the means *T*_*d*_*µ* (for drivers) and *T*_*p*_*µ* (for passengers), where *T*_*d*_ and *T*_*p*_ are the mutation target sizes for drivers and passengers, respectively. New driver mutations were randomly assigned to the cancer genes from the epistasis network; if a gene had already been hit by a previous driver mutation, all subsequent driver mutations in that gene were treated as neutral.

To study the effect of epistasis in cancer genome evolution, we simulated 5 scenarios represented by the following epistasis networks: (i) no epistasis, represented as a “network” of 10 disconnected genes; (ii) 2 × 5-node star networks, with positive epistasis between the central and peripheral genes of each star; (iii) 2 × 5-node star networks as in (ii), adding negative epistasis between the central genes of each star; (iv) 2 × 5-node clique networks, with positive epistasis among all gene pairs from the same clique, and negative epistasis affecting half of all possible gene pairs from different cliques; and (v) 3 × 5-node “negative clique” network, with negative epistasis among all gene pairs from the same clique, and positive epistasis affecting 10% of all possible gene pairs from different cliques (this percentage was chosen to achieve a positive-to-negative interaction ratio that inversely mirrored the synergistic clique scenario).

Simulations started from a clonal population of size *K* with one driver mutation in a randomly selected gene and no passengers. We used the following parameter values: *K* = 1000, *µ* = 10^−8^, *T*_*d*_ = 700, *T*_*p*_ = 5 × 10^6^, *s*_*p*_ = 10^−3^, *ϵ* = 5. To facilitate comparisons, the parameter *s*_*d*_ was separately set in each scenario, such that the average fitness effect when considering epistasis was the same (⟨*s*_*d*_⟩ = 0.1). Thus, in the disconnected and star networks (scenarios i-iii), we set *s*_*d*_ = ⟨*s*_*d*_⟩ = 0.1, in the clique network (scenario iv) *s*_*d*_ = 2⟨*s*_*d*_⟩/(1 + *ϵ*) ≈ 0.03, and in the negative clique network (scenario v) *s*_*d*_ = 3⟨*s*_*d*_⟩/*ϵ* = 0.06. Simulations ended when the cell population doubled its initial size, became extinct, or after 30,000 generations. The final composition of those populations that doubled the initial size (representing tumor progression) was used to investigate differentiation into mutation-based subtypes by running a principal component analysis on the mutation composition matrix that represented the most frequent genotypes of 500 independently evolving populations.

## Supplemental Information

**Supplementary Figure 1** (related to Figure 1d): Comparison of *Coselens* and SELECT methods for the detection of conditionally selected gene pairs, separately showing the overlap of both methods for positive and negative epistasis. CNA: copy number alterations.

**Supplementary Figure 2** (related to Figure 4): Epistasis networks for cancer types not shown in Figure 4. Nodes represent cancer genes involved in conditional selection; edges denote conditionally selected gene pairs. Cancer genes that are not involved in conditional selection are not represented, although they sometimes harbor a sizeable fraction of driver substitutions. Node size is proportional to the frequency of driver substitutions in a gene; edge width is proportional to the effect size; edge color represents the sign of epistasis; and arrows indicate the degree of symmetry (fully unidirectional arrows indicate lack of statistical significance and/or insufficient sample size for testing the reverse interaction).

**Supplementary Figure 3:** Kaplan-Meier curves showing association between conditional selection at the gene pair level and differential survival. The p-value indicated by *p*all corresponds to the log-rank test for the null hypothesis that there are no differences among the three survival curves; *p*1 and *p*2 correspond to pairwise comparisons between the double mutant and each of the single mutants (log-rank test).

**Supplementary Table 1:** Genes under significant levels of positive selection in at least one cancer type, used as “query genes” in this study. For each gene, the selection parameters for missense, nonsense, and indels, and the estimated number of driver mutations per tumor are shown. The table also indicates if a query gene was additionally considered as a “split gene”.

**Supplementary Table 2:** Gene pairs under significant conditional selection in at least one cancer type, sorted by effect size. For each pair, the table shows the mean number of driver substitutions per tumor in the query gene in the presence (Nd.with) and absence (Nd.without) of coding mutations in the split gene, the magnitude of conditional selection (DeltaNd), the dependence index (theta), the epistasis class, and the q-values for the null hypotheses Nd.with = 0, Nd.without = 0, and DeltaNd = 0.

**Supplementary Table 3:** Gene pairs that display mutual conditional selection, sorted by asymmetry. For each pair, the table shows the dependence indices in both directions (theta_AB and theta_BA), the asymmetry, the sign of epistasis, and the mean relative difference in variant allele frequency.

**Supplementary Table 4:** Analysis of conditional selection within cancer subtypes for breast cancer (BRCA), colorectal cancer (COAD), endometrial cancer (UCEC), and lower grade glioma (LGG). Statistical significance levels are coded as n.s. (not significant), * (p<0.05),** (p<0.01), and *** (p<0.01).

