## Supplementary figures and images for "Pervasive conditional selection of driver mutations and modular epistasis networks in cancer"

### Supplementary Figure S1

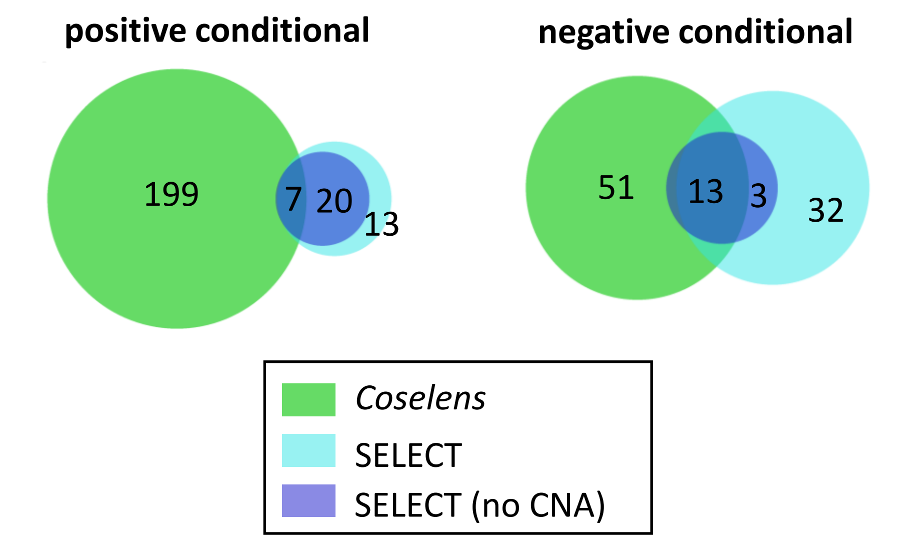

### Supplementary Figure S2

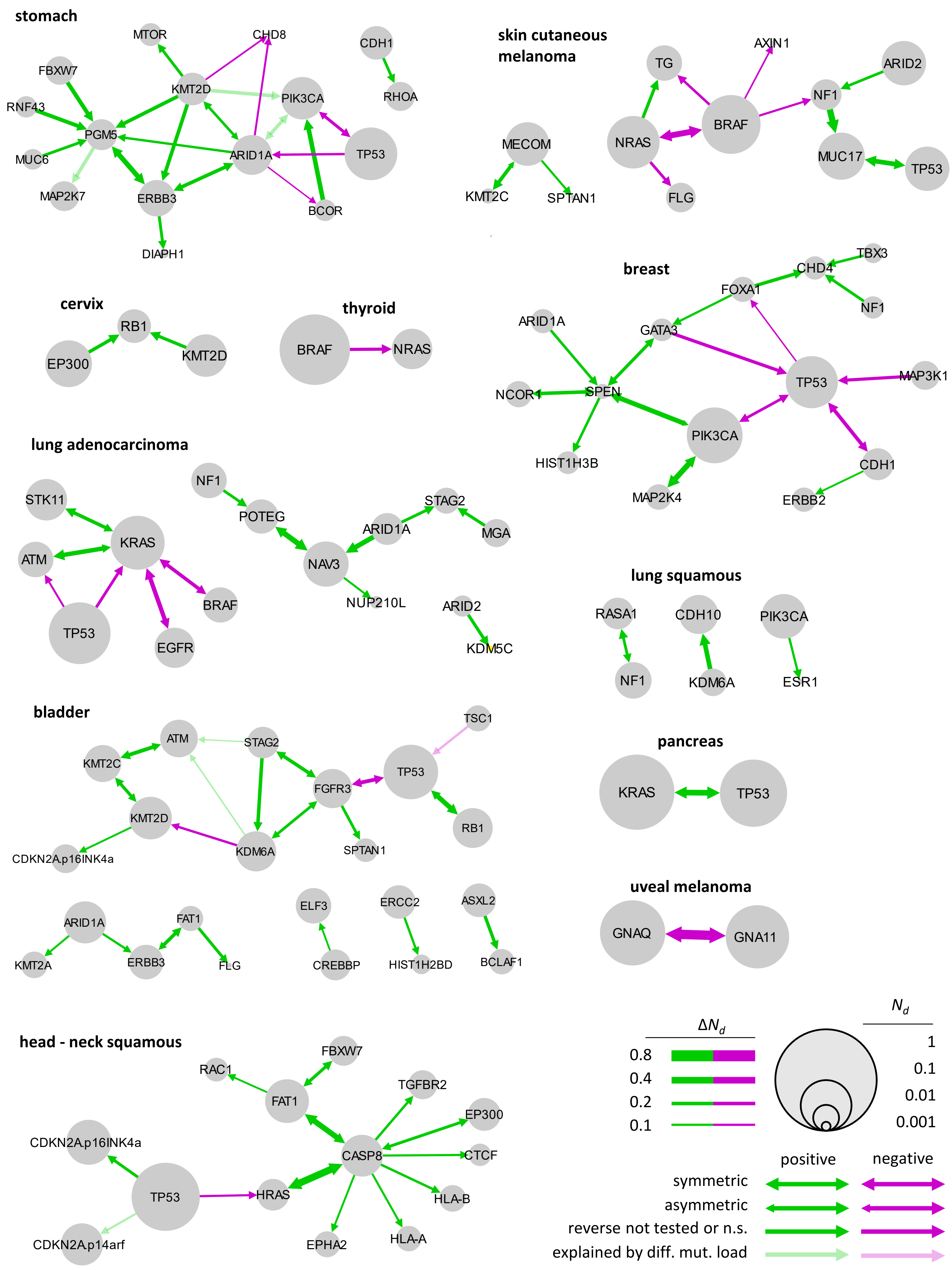

### Supplementary Figure S3

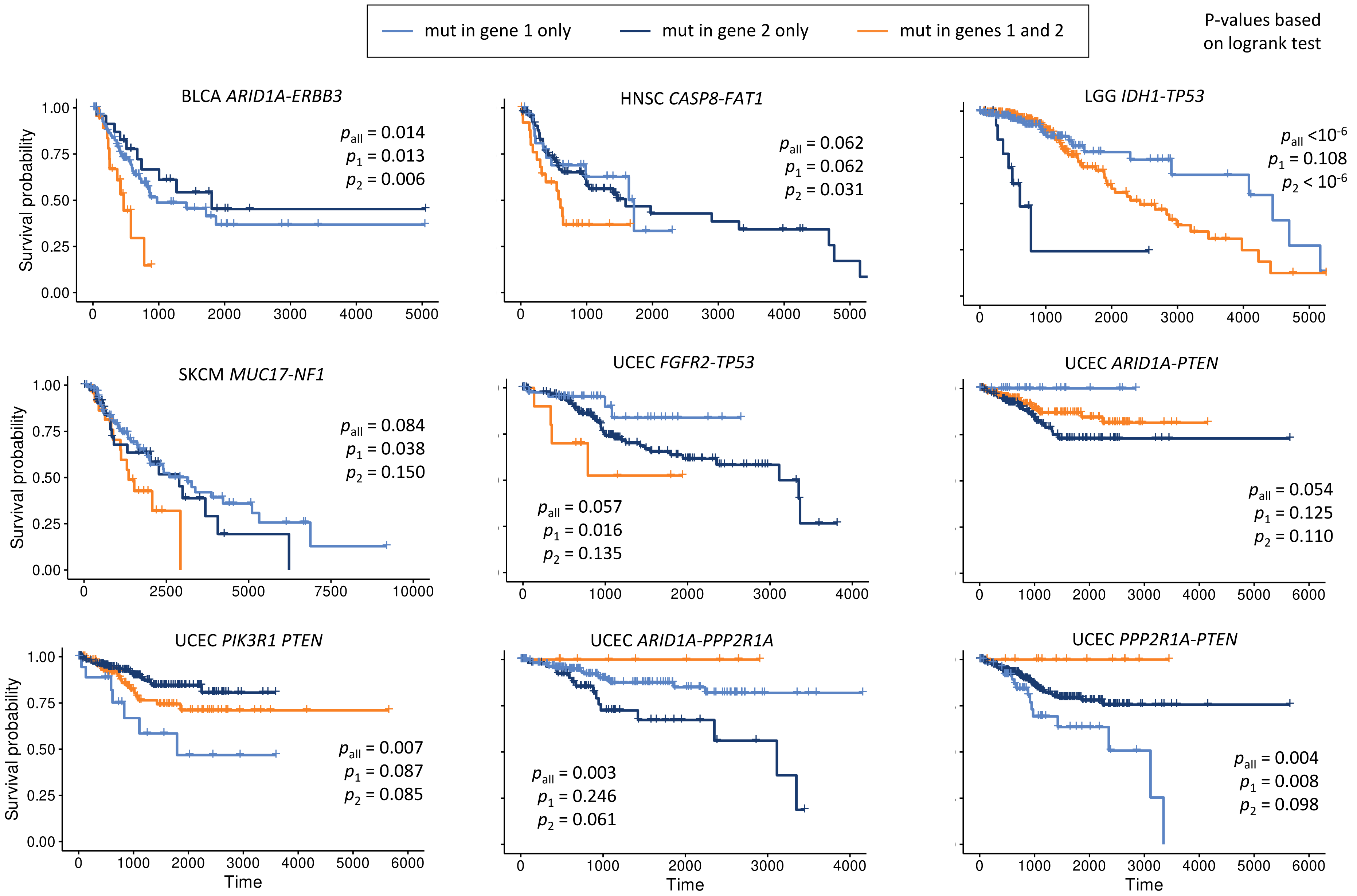
